# Reticular Dysgenesis-associated Adenylate Kinase 2 deficiency causes failure of myelopoiesis through disordered purine metabolism

**DOI:** 10.1101/2021.07.05.450633

**Authors:** Wenqing Wang, Andrew DeVilbiss, Martin Arreola, Thomas Mathews, Zhiyu Zhao, Misty Martin-Sandoval, Giorgia Benegiamo, Avni Awani, Ludger Goeminne, Daniel Dever, Yusuke Nakauchi, Mara Pavel-Dinu, Waleed Al-Herz, Luigi Noratangelo, Matthew H. Porteus, Johan Auwerx, Sean J. Morrison, Katja G. Weinacht

## Abstract

Reticular Dysgenesis is a particularly grave form of severe combined immunodeficiency that affects the adaptive and innate immune system. Patients suffer from congenital neutropenia, lymphopenia, and deafness. The disease is caused by biallelic loss of function in mitochondrial Adenylate Kinase 2 (AK2). AK2 mediates the phosphorylation of AMP to ADP, as substrate for ATP synthesis. Accordingly, declining oxidative phosphorylation has been postulated as the driver of disease pathology. The mechanistic basis, however, remains incompletely understood. Single cell RNA-sequencing of patient bone marrow cells implicated altered RNA catabolism and ribonucleoprotein synthesis in the pathogenesis of Reticular Dysgenesis. To investigate these findings, we developed a disease model based on CRISPR-mediated disruption of the *AK2* gene in primary human hematopoietic stem cells. We found that AK2-deficient myeloid progenitor cells not only have compromised mitochondrial energy metabolism and increased AMP levels, but also NAD^+^ and aspartate depletion, metabolites that rely on TCA-cycle activity for regeneration and synthesis. Furthermore, AK2-deficient cells exhibited strikingly increased levels of the purine nucleotide precursor IMP, decreased cellular RNA content, ribosome subunit expression, protein synthesis and a profoundly hypo-proliferative phenotype. The rise in IMP levels stemmed from increased AMP deamination. Pharmacologic inhibition of AMP deaminase normalized IMP levels in AK2-deficient cells, but further aggravated the disease phenotype, pointing to AMP catabolism as a possible metabolic adaptation to mitigate AMP-mediated toxicity. Inducing an adenosine disequilibrium in control cells produced a similar myeloid maturation defect.

This study shows that AK2 deficiency globally curtailed mitochondrial metabolism resulting in NAD^+^ and aspartate deficiency and disordered purine metabolism. AMP accumulation and its detrimental effects on ribonucleotide synthesis capacity may contribute to the failure of myelopoiesis in Reticular Dysgenesis.

## INTRODUCTION

The hematopoietic manifestations of Reticular Dysgenesis are defined by a combined maturation arrest in the myeloid and lymphoid lineages^1, 2^, while bone marrow cellularity is preserved and sometimes increased^3^. The disease is invariably fatal early in life unless hematopoietic and immune reconstitution are achieved by hematopoietic stem cell transplantation^4^. The predominance of opportunistic bacterial and fungal infections suggests that the neutropenia is the prevailing cause of death. Reticular Dysgenesis is caused by defects in the mitochondrial enzyme Adenylate Kinase 2^1, 2^. Interestingly, none of the classic mitochondriopathies caused by respiratory chain defects present with defects in myeloid or lymphoid development or function^5^.

Adenylate Kinases are a group of at least nine phosphotransferases that maintain adenine nucleotide homeostasis, i.e. the ratio between adenosine monophosphate (AMP), adenosine diphosphate ADP and (adenosine triphosphate) ATP in defined compartments of the cell^6–8^. Shuttle networks allow for the redistribution of AMP and ADP between spatially distinct locations of energy production and energy demand across the nucleotide-permeable outer mitochondrial membrane^6, 7, 9, 10^. AK2 is localized in the mitochondrial intermembrane space where it catalyzes the reversible reaction AMP+ATP↔2ADP. ADP is actively transported to the mitochondrial matrix by the ADP/ATP translocase where it is phosphorylated to ATP by ATP synthase. AK2 is ubiquitously expressed, while the expression of its cytosolic counterpart AK1 follows a tissue and cell type restricted pattern^2^. AK1 promotes the same enzymatic reaction as AK2, albeit at a lower rate (AK2 Km_(AMP)≤_10uM vs AK1 Km_(AMP)_=100-200mM)^6^. Because ADP is continuously removed from the intermembrane space while AMP and ATP are actively repleted, AK2 promotes the conversion of AMP and ATP to yield ADP. In the cytosol, the relative substrate concentrations are the reverse. Because ATP is consumed, AK1 mediates the conversion of ADP to ATP and AMP. Cytosolic AMP is shuttled back into the mitochondria for re-phosphorylation by AK2.

Although AK2 is ubiquitously expressed, non-hematopoietic tissues, with the exception of a subset of cells in the inner ear, seem unaffected by AK2 deficiency^1, 2^. This indicates that most cells do not depend on AK2 function. Within the hematopoietic system, platelets, erythrocytes, and some B cells also develop, at least to some extent, in patients with AK2 deficiency. Why AK2 function is dispensable in some hematopoietic and immune cells but not in others, is unclear. Interestingly, AK1 is not expressed in many hematopoietic and immune cells^2^. It has been speculated that the tissue specific phenotype of Reticular Dysgenesis stems from a lack of redundancy of AK1 and AK2 in cell types affected by Reticular Dysgenesis (e.g. neutrophils and T cells) and that AK1 may at least partially compensate for the lack of AK2 in cells in which both kinases are expressed (megakaryocytes, erythroid precursors, early pro B-cells) ^1, 2, 11, 12^.

Previous studies exploring the mechanism of Reticular Dysgenesis have largely focused on the compromise in OXPHOS metabolism caused by AK2 deficiency^12–15^. However, it remained unclear how this translates into the observed defect in proliferation and differentiation. Moreover, cells have a myriad of redundantly ADP-producing pathways that can fuel ATP synthesis and the specific cell types that are predominantly affected in Reticular Dysgenesis, i.e. neutrophils and T cells, are thought to rely heavily on glycolysis for ATP production, both at baseline and under stress^16–22^. Therefore, alternate drivers of disease pathology need to be considered. Progress has been hindered by lack of appropriate model systems to phenocopy the human disease. To understand the cellular consequences of AK2 deficiency on the human hematopoietic system, we performed single cell RNA-sequencing on the bone marrow of Reticular Dysgenesis patients and established a novel AK2 CRISPR-knock out model in primary human hematopoietic stem cells.

## MATERIAL AND METHODS

### Human subjects

All specimens from human subjects were obtained under informed consent in accordance with the Boston Children’s Hospital Institutional Review Board.

### Single cell RNA-sequencing

Bone marrow samples from two Reticular Dysgenesis patients were obtained in EDTA per institutional protocol. Red blood cell lysis was performed with ammonium chloride solution (StemCell Technologies 07850). Single cell RNA-seq libraries were constructed from FACS sorted CD45+ bone marrow cells, using Chromium Single Cell 3ʹ GEM, Library & Gel Bead Kit v3 (10X Genomics). Libraries were sequenced on Illumina NextSeq 500. Single cell RNA-seq data from bone marrow mononuclear cells of eight healthy human donors available through the Human Cell Atlas served as control^23^. Read alignment to the human genome hg19, cell barcode calling and UMI count were performed using Cellranger 3.0.0. Cell clustering and differential gene expression were performed using Seurat v3^24^.

### Cell culture

Human cord blood CD34^+^ hematopoietic stem and progenitor cells (HSPCs) were purchased (StemCell Technologies #78003) or acquired through the Binns Program for Cord Blood Research at Stanford University under informed consent. Human fetal liver CD34^+^ HSPCs were isolated from fetal liver tissues acquired through Advanced Bioscience Resources, Inc. For CD34+ HSPC *in vitro* expansion, cells were cultured in StemSpan SFEMII (StemCell Technologies 09655) supplemented with 100μg/mL SCF (StemCell Technologies 78062), 100μg/mL TPO (StemCell Technologies 78210.1), 100μg/mL FLT3L (StemCell Technologies 78009), 100μg/mL IL-6 (StemCell Technologies 78050), 35nM UM171 (StemCell Technologies 09655) and 0.75μM StemRegenin 1 (StemCell Technologies 72342), at 37°C, 5% O2 and 5% CO2.

For *in vitro* granulocyte differentiation, CD34^+^ HSPCs were cultured in Myelocult H5100 (StemCell Technologies 05150) supplemented with 100μg/mL hrSCF, 100μg/mL hrIL-3 (StemCell Technologies 78040), 10μg/mL hrG-CSF (NEUPOGEN, Amgen) and 1μM hydrocortisone (Sigma H0888) at 37°C and 5% CO2. Nicotinamide riboside (Elysium) was supplemented at 1-2mM. Aspartate (Sigma 11189) was supplemented at 10-20mM. AMPD inhibitor (Sigma 5.33642) was supplemented at 20μM.

For nucleoside deprivation and disequilibrium studies, CD34+ HSPCs were differentiated in MEM alpha, the base media of Myelocult H5100, with or without nucleosides (Thermo Fisher 32571-036 and 32561-037) with 10% dialyzed FBS (Thermo Fisher, A3382001), hrSCF, hrIL-3, hrG-CSF and hydrocortisone. Adenosine (Sigma A4036) or guanosine (Sigma G6264) was supplemented in MEM alpha without nucleosides at 40mg/L.

### rAAV6 production

rAAV6 was packaged using AAV vector plasmids and the rep/cap/helper plasmid pDGM6. The AAV vector plasmids were cloned in the pAAV-MCS plasmid containing ITRs from AAV serotype 2 (AAV2). Vectors contain an SFFV promoter, GFP or BFP reporter gene, BGH polyA, and left and right homology arms (400bp each) specific to *AK2^-/-^* or *AAVS1^-/^*^-^ CRISPR editing sites. pDGM6, a gift from David Russel, contains AAV6 cap, AAV2 rep, and adenovirus helper genes. rAAV6 particles were either produced by Vigene or as described with modifications^25^. Briefly, 293T cells grown to 70-80% confluency were transfected with 6 μg ITR-containing plasmid and 22 μg pDGM6 using 112 μg acidified PEI (Polysciences, 23966-1). 48-72 hours after transfection, cells were harvested and rAAV6 particles were purified using an AAVpro Purification Maxi kit (Takara, 6666) according to manufacturer’s protocol. rAAV6 titer was quantified by qRT-PCR following the previously described protocol^26^.

### CRISPR/Cas9 gene editing

CD34+ HSPCs were CRISPR edited at *AK2* or *AAVS1* loci, respectively, after 72-hour expansion in StemSpan. Synthetic sgRNAs were purchased from Synthego. sgRNA sequences targeting *AK2* or *AAVS1* locus are: *AK2*, UUCCUGUUCUCAGAUCACCG; *AAVS1*, GGGGCCACUAGGGACAGGAU. For 1 million CD34+ HSPCs, 8μg sgRNA and 15 μg Cas9 (IDT, 1081059) were mixed and incubated at room temperature for 15 minutes. RNP was delivered by electroporation using a Lonza 4D-nucleofector and P3 Primary Cell Kit (Lonza, V4XP-3012). Cells were transferred to fresh StemSpan media and corresponding (*AK2* or *AAVS1*) AAV vectors with GFP and BFP reporters were added to cell culture mix at an MOI of 50,000/cell. Cells were cultured for 3 days followed by FACS enrichment for lin-CD34+GFP+BFP+ cells.

### Flow cytometry and cell sorting

The following antibodies were used for flow cytometry and cell sorting of human bone marrow samples: CD117 PE (Biolgend 323408), CD16 PE-CF594 (BD 562293), CD33 PE-Cy7 (BD 333946), siglet-8 APC (Biolegend 347105), HLA-DR APC-R700 (BD 565127), CD14 APC-Cy7 (Biolgend 367108), CD11b BV605 (Biolgend 301332), CD3 BV650 (Biolgend 317324), CD19 BV650 (Biolgend 302238), CD56 BV650 (Biolgend 362532), CD34 BV711 (BD 740803), CD45 BV785 (Biolegend 304048), and CD15 BUV395 (BD 563872). The following antibodies were used for cell sorting of lin-CD34+GFP+BFP+ HSPCs after CRISPR editing: CD34 APC (Biolegend 343510), CD11b APC-Cy7 (Biolegend 301342), CD14 APC-Cy7 (Biolegend 367108), CD15 PE (Biolegend 323006), and CD16 PE (Biolegend 360704). The following antibodies were used for flow cytometry of myeloid differentiation and cell sorting of PMs, MCs and NPs: CD16 APC (Biolegend 302012), HLA-DR APC-Cy7 (Biolegend 307618), CD11b PE (Biolegend 301306) or BV711 (Biolegend 301344), CD117 PE-Cy7 (Biolegend 323408), and CD15 BV785 (Biolegend 323044). Cell sorting was performed on BD FACSAria II or FACSAria Fusion. Data analysis was done using Flowjo v10.

### Western blot

Primary antibodies used for Western blot of β-actin and AK2 proteins were: anti-actin (Santa Cruz Biotechnology sc-47778, 1:1000), anti-AK2 (ProteinTech 11014-1-AP, 1:200), anti-NDUFA9 (Abcam ab14713, 1:5000), anti-SHDA (Abcam, ab14715, 1:5000), anti UQCRCI (Abcam, ab110252, 1:5000), anti-ATP5A (Abcam, ab14748, 1:2000), anti-AMPK (Cell Signaling Technology, 2532s, 1:50), anti-pAMPK (Cell Signaling Technology, 2535s, 1:50). Secondary antibodies were: m-IgGκ BP-HRP (Santa Cruz Biotechnology sc-516102, 1:5000), mouse anti-rabbit IgG-HRP (Santa Cruz Biotechnology sc-2357, 1:5000).

### Electron transport chain supercomplex detection

Electron transport chain supercomplex detection was performed as previously described^27^. In brief, mitochondria were extracted from 10-20 million of unmanipulated HSPCs, PMs, MCs and NPs, and *AK2^-/-^* and *AAVS1^-/-^* MCs. For each sample, ∼15 μg mitochondria were run on a blue native PAGE gel and western transferred for immunoblotting. For detection of ETC supercomplexes anti-oxphos cocktail (Invitrogen, 457999) was used.

### Metabolomics

Cell sorting for metabolomics analysis was carried out as described^28, 29^. Briefly, cells were sorted on a FACSAria Fusion sorter running with a sheath fluid of 0.5X PBS, and a 70-μm nozzle. For each sample, 10,000 cells were sorted directly into 40μL acetonitrile. Samples were vortexed for 1 min to lyse the cells, then centrifuged at 17,000g for 15 min at 4 °C. Supernatant was transferred to a fresh tube and stored at −80 °C. For quantification of reduced and oxidized glutathione, 10,000 cells were sorted directly into 80% methanol with 0.1% formic acid. Samples were similarly extracted by vortexing. The supernatant was transferred to a fresh tube, dried down and reconstituted in 50 uL water with 0.1% formic acid. Metabolites were quantified on a Quadrupole-Orbitrap mass spectrometer (Thermo Fisher). Data analysis was performed using Omics Data Analyzer (https://git.biohpc.swmed.edu/CRI/ODA)^29^.

### RNA-sequencing

RNA was extracted from sorted promyelocytes, myelocytes and neutrophils using a Qiagen RNeasy Plus Micro kit. RNA-seq library was constructed with a Takara SMARTseq v4 kit and sequenced on Illumina NovaSeq. Reads mapping to the human genome hg38, transcript quantification and TPM calculation were performed using Kallisto v0.46.0^30^. Differential gene expression was analyzed by DESeq2^31^.

### Seahorse mitochondrial stress test

Real-time cell metabolic analysis was performed on a Seahorse XFe96 Analyzer (Agilent) using a XF Cell Mito Stress Test kit (Agilent 103015-100) according to the manufacturer’s protocol. In brief, a 96-well Seahorse cell culture plate was coated with 0.1 mg/mL Poly-D-Lysine (Advanced Biomatrix 5174). Sorted promyelocytes, myelocytes and neutrophils were pelleted and resuspended in Seahorse XF RPMI media (Agilent 103576-100). 80,000-100,000 cells were plated in each well. The cell culture plate was gently centrifuged to facilitate settling. Each cell type/treatment condition was plated at least in triplicate. Oligomycin (final concentration 1.5 uM), FCCP, (final centration 1 uM) and rotenone/antimycin A (final concentrations 0.5 uM each) were injected into the media sequentially during the assay, and oxygen consumption rate and extracellular acidification rate were measured before and after each injection.

### Mitochondrial copy number assay

Genomic DNA was extracted using a Qiagen DNeasy Blood and Tissue kit. qPCR quantification of mitochondrial copy number was performed as described^32^. Primers and probes for nuclear B2M and mitochondrial MT-ND1 genes were purchased from IDT. B2M-F, CCAGCAGAGAATGGAAAGTCAA; B2M-R, TCTCTCTCCATTCTTCAGTAAGTCAACT; B2M-Probe (FAM), ATGTGTCTGGGTTTCATCCATCCGACA; MT-ND1-F, CCCTAAAACCCGCCACATCT; MT-ND1-R, GAGCGATGGTGAGAGCTAAGGT; MT-ND1-Probe (FAM), CCATCACCCTCTACATCACCGCCC.

### Mitochondrial membrane potential

To quantify mitochondrial membrane potential, cells were stained with 20 nM TMRM (Thermo Fisher T668) in PBS, incubated at 37°C and 5% CO2 for 1 hour, and analyzed by flow cytometry.

### RNA content

RNA content was quantified by Pyronin Y staining. 500,000 cells were plated in 1 mL fresh Myelocult media. Hoechst 33324 (ThermoFisher H3570) was added to the media at the concentration of 5 μg/mL, and cells were incubated at 37 °C and 5% CO2 for 30 minutes. Pyronin Y (Sigma P9172) was then added at a concentration of 0.5 μg/mL. Cells were incubated for an additional 45 minutes. Subsequently, cells were stained with surface marker antibodies and analyzed by flow cytometry.

### Nascent peptide synthesis

The Click-iT™ Plus OPP Alexa Fluor™ 594 Protein Synthesis Assay Kit (Thermo Fisher C10457) was used to quantify nascent peptide synthesis. 500,000 cells were plated in 1 mL fresh Myelocult media supplemented with 20 μM OP-puromycin (OPP) and incubated for 1 hour at 37 °C and 5% CO2. Subsequently, cells were collected for cell surface marker staining, fixation with 1% paraformaldehyde, permeabilization with 0.1% saponin, and OPP staining according to the manufacturer’s protocol. Cells were then analyzed by flow cytometry.

### Statistical Analysis

p values were determined by two-way ANOVA with Tukey’s post-test, paired or unpaired Student’s t test using R/Bioconductor. Error bars represent standard deviations. Differences were considered statistically significant when p < 0.05 (* p < 0.05, ** p < 0.01, *** p < 0.001, **** p < 0.0001, ns = not significant).

### Data Availability

Single cell RNA-seq data was deposited in GEO with accession number GSE179346. RNA-seq was deposited in GEO with accession number GSE179320.

## RESULTS

### Single cell RNA sequencing of Reticular Dysgenesis patient bone marrow

To glean insights into the mechanistic basis of Reticular Dysgenesis (RD), we performed single cell RNA sequencing (scRNA-seq) on bone marrow samples of two previously reported patients with biallelic c.542G>A mutation^4, 11, 12^. Bone marrow scRNA-seq data from 8 healthy donors served as control^23^. Unsupervised clustering using Seurat identified 22 individual hematopoietic progenitor and mature hematopoietic cell populations (Fig 1A, S1). RD patient bone marrow comprised a significantly larger fraction of immature cells, i.e. hematopoietic stem cells (HSCs), common myeloid progenitor cells (CMPs), common lymphoid progenitor cells (CLPs), pro-B cells (ProBs), megakaryocyte-erythroid progenitors (MEPs) and early erythroid progenitors (ERPs), while mature populations across myeloid and lymphoid lineages were significantly decreased (Fig 1B, C, S2). Erythroid development was the least affected in the RD samples with relatively normal maturation into late erythroid progenitors (Fig 1B, C, S2), possibly due to the compensatory activity of AK1, which is expressed in the erythroid lineage, but not in hematopoietic stem and progenitor cells (HSPCs), myeloid or lymphoid populations (Fig S3). Capturing fully mature granulocytes by the droplet-based 10X scRNA-seq platform remains challenging^33^, due to high levels of RNases/proteinases in this cell type. Therefore, neutrophil populations were neither captured in patient nor control samples. The myeloid differentiation defect demonstrated by scRNA-seq was also observed by flow cytometry (Fig S4). In the myeloid compartment (CD45^+^ CD3^-^ CD19^-^ CD56^-^), RD patients showed significantly lower commitment to the granulocytic lineage (HLA-DR^-^), and an expansion of promyelocytes (CD117^+^) combined with a decrease in the frequency of cells at more mature granulocyte committed myeloid stages (CD15^+^ CD11b^+^ CD16^+^).

**Figure 1.**
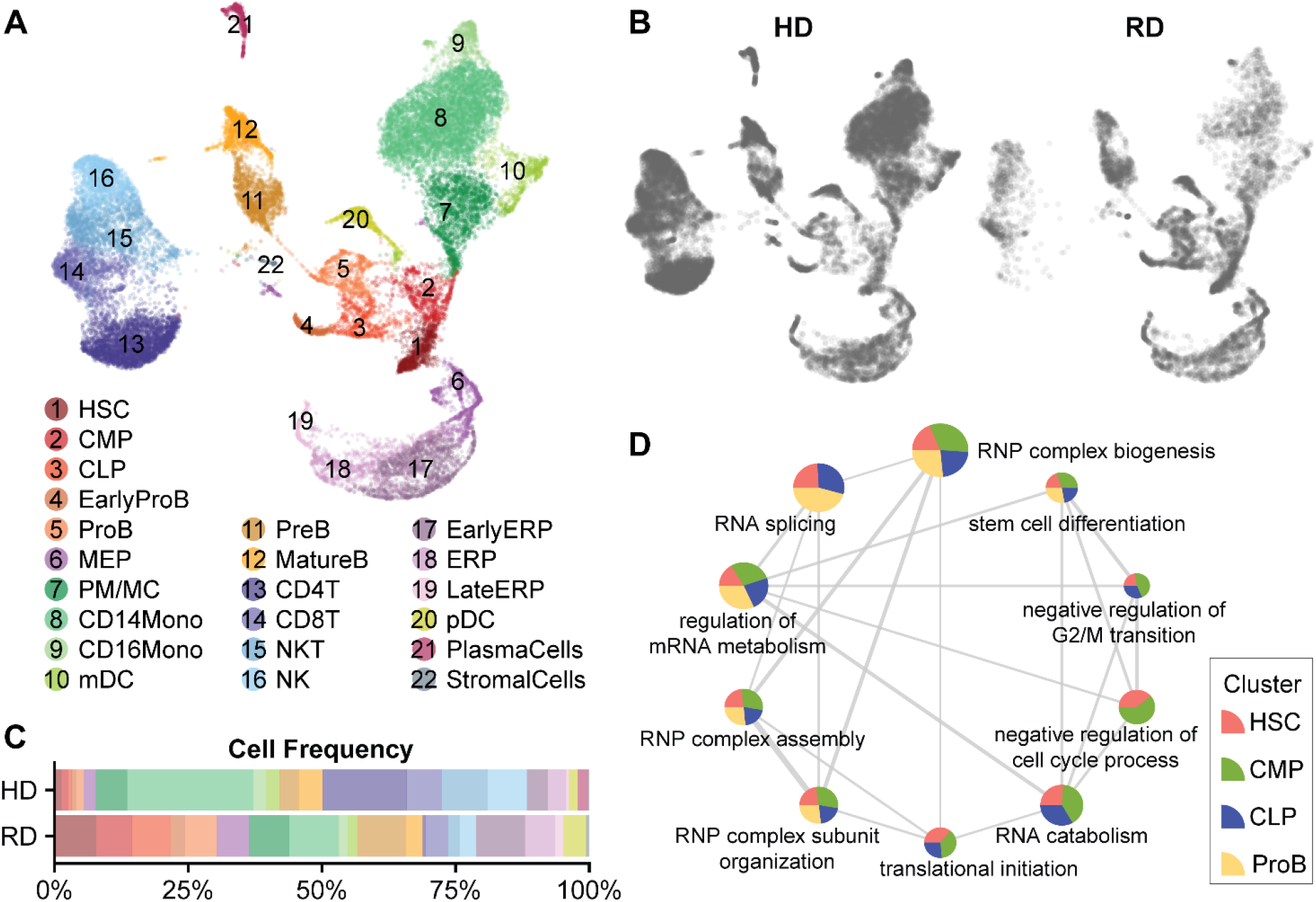
Single cell RNA-seq analysis of Reticular Dysgenesis patient bone marrow reveals maturation arrest across multiple lineages and defects in RNA metabolism and ribosome biogenesis. (A) UMAP of bone marrow cells from 2 Reticular Dysgenesis patients (RD) and 8 healthy donors (HD) identified 22 clusters. HSC, hematopoietic stem cells; CMP, common myeloid progenitor cells; CLP, common lymphoid progenitor cells; B, B cells; mDC, myeloid dendritic cells; MEP, megakaryocyte–erythroid progenitor cells; PM/MC, promyelocytes/myelocytes; NK, natural killer cells; NKT, natural killer T cells; ERP, erythroid progenitor cells; pDC, plasmacytoid dendritic cells. (B, C) Feature plots representing distribution of RD and HD bone marrow cells across all clusters. RD samples showed a higher frequency in cells of HSPC clusters (HSC, CMP, CLP, EarlyProB, ProB and MEP), and lower frequency in cells of mature myeloid and lymphoid clusters. (D) GO enrichment map of down regulated differentially expressed genes (DEGs) in HD vs RD cells, plotted with ClusterProfiler^1^. Pie size of each node is proportional to the number of DEGs in each GO gene set. Edge width is proportional to number of overlapping genes between gene sets. Multiple pathways related to RNA metabolism, ribosomal biogenesis and cell cycle progression are enriched in HSPC clusters (HSC, CMP, CLP and ProB) of RD patient bone marrow cells. All enriched pathways are downregulated in RD cells compared to controls.

A differential gene expression and gene set enrichment analysis comparing individual HSPC clusters (HSC, CMP, CLP and pro-B cells) in RD patients and healthy donors, revealed a distinct set of pathways that were downregulated in RD patients, with high congruency between the different HSPC cell types queried (Fig 1D). The top 10 differentially regulated pathways in HSPC clusters included pathways involved in RNA metabolism, ribonucleoprotein (RNP) complex biogenesis and proliferation/cell cycle progression, which were all significantly downregulated in RD patients (Fig 1D). These findings led us to hypothesize that defective ribonucleotide metabolism underlies the proliferation and differentiation arrest leading to neutropenia and immunodeficiency in RD.

### A Cell-traceable CRISPR *AK2* Deletion Model

A number of strategies have been explored to model RD. *Ak2*-deficient mice, including germline deficiency^34^ and lineage specific, hematopoietic deletion with *Vav*-Cre are embryonically lethal^35^. Prior studies using *ak2*^-/-^ zebrafish^12^, human cell *AK2* shRNA knock-down^36^ and our work in *AK2*-deficient human iPSCs^12^ advanced our understanding of RD, but none of these models fully recapitulated definitive human hematopoiesis. To overcome prior limitations and precisely recapitulate the failure of human myelopoiesis in culture, we designed a CRISPR construct to disable *AK2* in primary human CD34^+^ HSPCs. Because only complete absence of *AK2* expression reproduces the RD phenotype, disruption of both alleles is required. We combined CRISPR/Cas9 gene editing with adeno-associated viral vector delivery of two homologous donors containing GFP and BFP reporters^25, 37^ (Fig 2A), to disrupt the *AK2* gene at the catalytic LID domain. This strategy allows selection of biallelically edited cells (GFP+ BFP+) by flow cytometry (Fig 2B). *AK2*-edited GFP^+^ BFP^+^ HSPCs (*AK2-/*-) show complete absence of the AK2 protein (Fig 2C). Cells edited at the safe harbor locus, *AAVS1*^38^, were used as a negative control (*AAVS1*^-/-^). To mimic the RD-specific neutropenia phenotype, *AK2^-/-^* and *AAVS1^-/-^* HSPCs were differentiated *in vitro* along the granulocytic lineage (Fig 2D). Consistent with the presentation in RD patients (Fig S4), *AK2^-/-^* HSPCs showed decreased commitment to the HLA-DR^-^ granulocytic lineage (Fig 2E, F), arrested differentiation at the promyelocyte (PM) stage (CD15^-^ CD117^+^), and failure to further mature into myelocytes (MCs, CD15^+^ CD11b^+^ CD16^-^) and neutrophils (NPs, CD15^+^ CD11b^+^ CD16^+^) (Fig 2E, G). Strikingly, overall proliferative capacity of *AK2^-/-^* myeloid progenitors was severely compromised (Fig 2H), resulting in a significantly lower yield of MCs and NPs (Fig 2I). These results demonstrate that the biallelic AK2 deficiency recapitulates the failure of myelopoiesis in RD with high fidelity.

**Figure 2.**
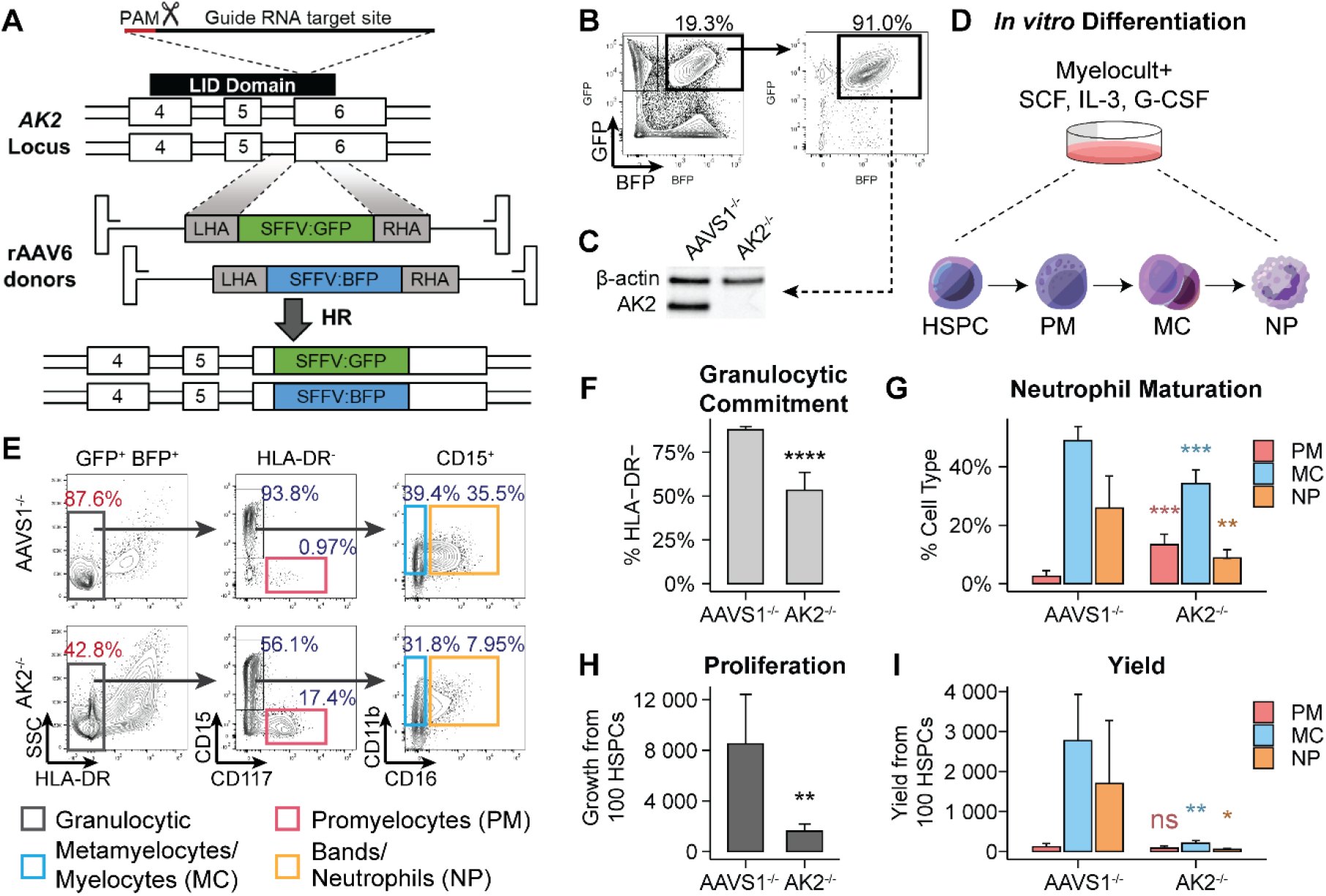
A biallelic AK2 knock out CRISPR/Cas9 model in human HSPCs recapitulates defective myelopoiesis in RD. (A) Design of biallelic AK2 knock out CRISPR/Cas9 model. *AK2* guide RNA (gRNA) targets the LID domain. Two rAAV6 donors, containing left and right homologous arms (LHA and RHA) flanking the *GFP* and *BFP* reporters, respectively, are used as repair templates to introduce *AK2* loss-of-function mutations following CRISRP/Cas9 DNA cutting at the gRNA targeting site. (B) Following CRISPR/Cas9 gene editing, cells with biallelic *AK2* disruption are selected by GFP+ BFP+ sorting 3 days after CRISPR/Cas9 gene editing. >90% of cells remain GFP+ BFP+ at the end of the *in vitro* culture. Successfully edited HSPCs are referred to *AK2^-/-^* and *AAVS1^-/-^* cells. (C) Western blot confirms the absence of AK2 protein expression in the GFP+ BFP+ fraction of *AK2* CRISPR edited HSPCs. (D) *AAVS1^-/-^* and *AK2^-/-^* HSPCs are differentiated *in vitro* along the granulocytic lineage in Myelocult™ media for 7 days. (E) Representative flow cytometry plots of *AAVS1^-/-^* and *AK2^-/-^* cells following neutrophil differentiation. (F) Percentages of HLA-DR-cells in *AAVS1^-/-^* and *AK2^-/-^* cells. (G) Percentages of PMs, MCs and NPs within the HLA-DR-granulocytic lineage in *AAVS1^-/-^* and *AK2^-/-^* cells. (H) Proliferation of *AAVS1^-/-^* vs *AK2^-/-^* cells during neutrophil differentiation. (I) Yield of PMs, MCs and NPs was determined by the number of cells at the end of culture normalized to 100 HSPCs at the beginning of the culture. (F-I) showed data of 8 biological replicates. Error bars represent standard deviation.

### Cellular energy demand peaks at the promyelocyte stage of human myelopoiesis

Quiescent long-term HSCs reside in a hypoxic microenvironment and retain a low metabolic profile^39, 40^. Short-term HSCs and multipotent progenitor cells (MPPs) undergo a shift towards mitochondrial oxidative phosphorylation (OXPHOS)^41, 42^. Mature neutrophils return to glycolysis to support ATP production and exhibit a low rate of OXPHOS^43, 44^. Regulation of energy metabolism at the stages in between remains incompletely understood. To explore why *AK2*-deficient myelopoiesis arrests at the PM stage, we investigated how patterns of energy production change during development. Using a Seahorse extracellular flux analyzer, we quantified mitochondrial respiration and glycolysis rate in unmanipulated HSPCs, PMs, MCs and NPs derived from our *in vitro* model (Fig 3A, B). Baseline mitochondrial respiration, mitochondrial copy number, protein expression of electron transport chain (ETC) complexes, specifically the NADH-dependent ETC complexes I (NDUFA9) and III (UQCRC1) as well as ETC super-complexes (SC), all peaked at the PM stage and then steadily declined during terminal differentiation into NPs (Fig 3C, F-I). Maximal respiration and spare respiratory capacity were highest in HSCs. (Fig 3D, E). Interestingly, the rate of glycolysis also peaked at the PM stage and then declined (Fig 3I), suggesting that energy metabolism globally peaks at the promyelocyte stage (Fig 3J).

**Figure 3.**
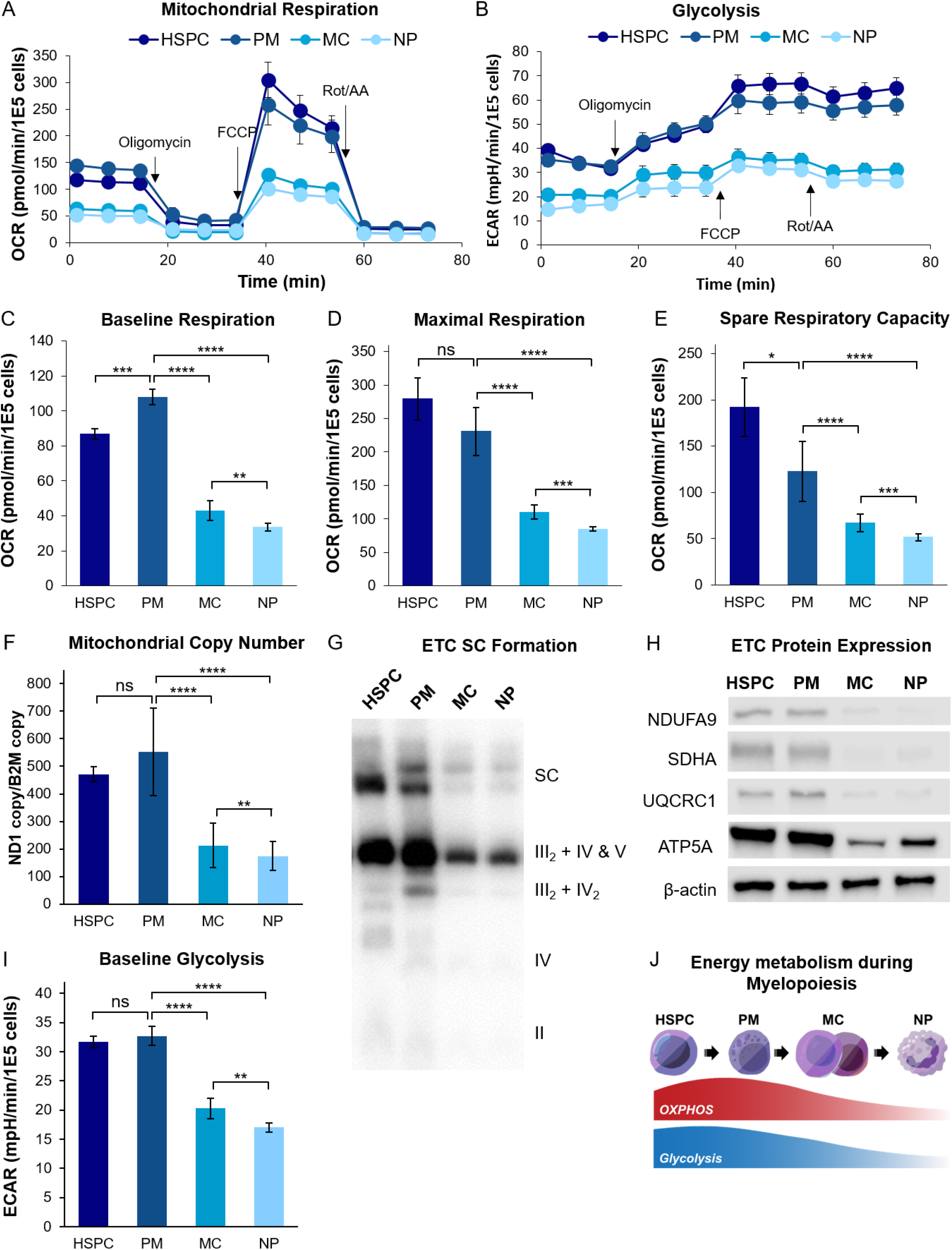
Energy metabolism peaks at the promyelocyte stage of myelopoiesis. (A, B) Seahorse mito stress test assay measuring oxygen consumption rate (OCR) and extracellular acidification rate (ECAR) in unmanipulated HSPCs, PMs, MCs and NPs. Oligomycin, FCCP and rotenone/antimycin A were sequentially injected into cell culture wells. OCR and ECAR were measured before and after each drug injection. 3-5 replicates were plated for each cell type. (C) Quantification of baseline OCR, (D) maximal OCR, (E) spare respiratory capacity (maximal OCR - baseline OCR), (F) Mitochondrial copy numbers of each cell type from 3 biological replicates. (G) Quantification of electron transport chain (ETC) complexes and super-complexes (SC) in mitochondrial extract of each cell type, visualized by blue native gel electrophoresis followed by western blotting using complex I-V specific antibodies. (H) Representative western blot of electron transport chain complex proteins in each cell type. NDUFA9, complex I; SDHA, complex II; UQCRC1, complex III; ATP5A, complex V. β-actin is loading control. (I) baseline ECAR for each cell type. Error bars indicate standard deviation. (J) Scheme of dynamic OXPHOS and glycolysis demand during myelopoiesis.

### AK2 deficiency leads to a mild reduction in mitochondrial ATP synthesis

To further investigate the metabolic defects conferred by AK2 deficiency, we focused on the energetically most active promyelocyte stage, which is the developmental stage where proliferation peaks^18^. To approximate substrate turnover over time, we analyzed mitochondrial respiration and glycolysis in *AK2^-/-^* and *AAVS1^-/-^* cells using an extracellular flux analyzer (Fig 4A, 4B). *AK2^-/-^* PMs exhibited a lower baseline respiration (Fig 4C), but normal maximal respiration and normal (with a trend towards increased) spare respiratory capacity upon pharmacologic uncoupling (Fig 4D, E). The rate of glycolysis was slightly increased in *AK2^-/-^* PMs (Fig 4F). Furthermore, *AK2^-/-^* PMs and MCs showed no change in mitochondrial copy number (Fig 4G) and AK2 deficiency did not impact the expression of ETC complexes and super-complexes as compared to control *AAVS1^-/-^* cells (Fig 4H). Interestingly, the mitochondrial membrane potential was slightly higher in *AK2^-/-^* PMs as compared to control PMs (Fig 4I). The mitochondrial membrane potential is maintained by the activity of respiratory chain complexes I and III and drives the activity of the mitochondrial ATP synthase. Lower baseline respiration and higher membrane potential in *AK2*-deficient cells may be a consequence of limited ADP availability consistent with the primary function of AK2. Normal mitochondrial spare respiratory capacity, a measure of electron transport chain capacity and normal expression of ETC complexes and supercomplexes in AK2-deficient cells make compromised respiratory chain integrity unlikely to drive the pathogenesis of RD. The increase in ATP synthesis from glycolysis points to a compensatory adaptation to meet cellular energy demand. Overall, these data reveal no evidence of a catastrophic loss of ATP synthesis, necessitating broader investigations into the consequences of AK2 deficiency to understand the defect in proliferation and differentiation.

**Figure 4.**
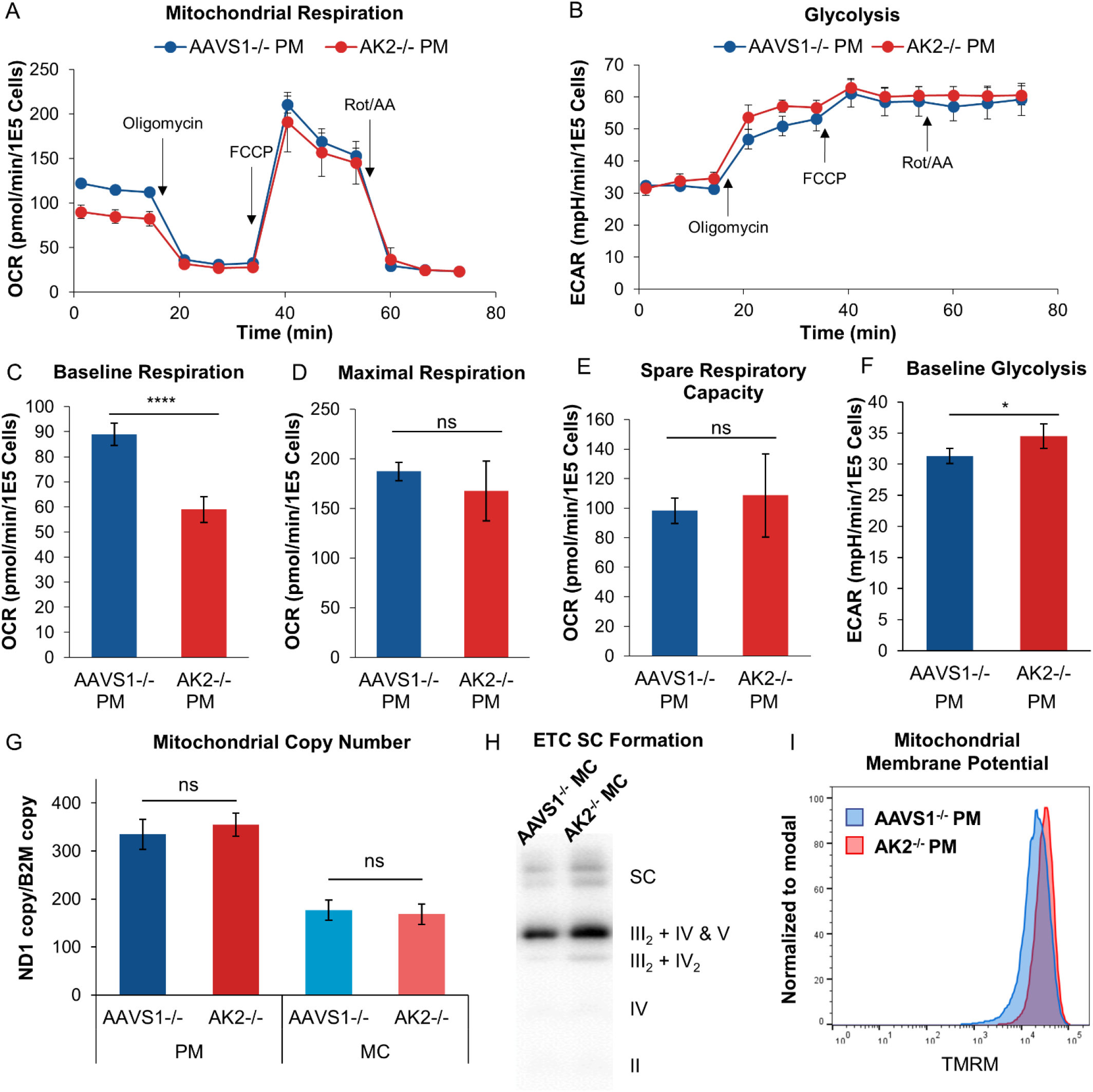
AK2-deficiency causes mild reduction in ATP synthesis without compromising respiratory chain integrity. (A, B) Seahorse mito stress test assay measuring OCR and ECAR of *AAVS1^-/-^* and *AK2^-/-^* PMs. Oligomycin, FCCP and rotenone/antimycin A were sequentially injected into cell culture wells, OCR and ECAR were measured before and after each drug injection. 3-5 replicates were plated for *AAVS1^-/-^* and *AK2^-/-^* PMs each. (C-F) Quantification of baseline OCR, maximal OCR, spare respiratory capacity, and baseline ECAR for *AAVS1^-/-^* and *AK2^-/-^* PMs. Error bars indicate standard deviation. (G) Mitochondrial copy numbers of *AAVS1^-/-^* and *AK2^-/-^* PMs and MCs from 3 biological replicates. (H) Quantification of electron transport chain (ETC) complexes and super-complexes (SC) in mitochondrial extract of *AAVS1^-/-^* and *AK2^-/-^* cells, visualized by blue native gel electrophoresis followed by western blotting using complex I-V specific antibodies. (I) Mitochondrial membrane potential of *AK2^-/-^* vs *AAVS1^-/-^* PMs, measured by TMRM staining. Representative flow cytometry results from 3 independent experiments were shown.

### AK2 deficiency causes a broad decline in mitochondrial biosynthetic activity

To understand the global metabolic impact of AK2 deficiency, we performed metabolomic profiling using LC-MS/MS^28, 29^ in *AK2^-/-^* and *AAVS1^-/-^* PMs, MCs and NPs. As expected, AK2 deficiency increased AMP levels, and AMP/ADP and AMP/ATP ratios at the MC and NP stages (Fig 5A) while ADP and ATP steady state levels were not significantly changed as compared to *AAVS1^-/-^* controls (Fig S6A). This suggests that adenine nucleotide homeostasis in AK2-deficient cells is primarily disrupted by an accumulation of AMP, while ADP and ATP levels are close to normal, possibly by curtailing ATP flux or tapping into alternate pathways for ADP and ATP generation. Unexpectedly, we observed a profound decrease in nicotinamide adenine dinucleotide (NAD^+^) levels, which led to a significant increase in NADH/NAD^+^ ratios, while NADH levels remained unchanged (Fig 5B, S6B). The increase in NADH/NAD^+^ ratio suggests a relative excess of reducing equivalents in *AK2^-/-^* cells, which is corroborated by an increase in the ratio of reduced to oxidized glutathione (GSH/GSSG) (Fig 5B), the primary mitochondrial redox couple. Notably, AMP accumulation and redox imbalance progressively worsened as cells matured from PMs to NPs.

**Figure 5.**
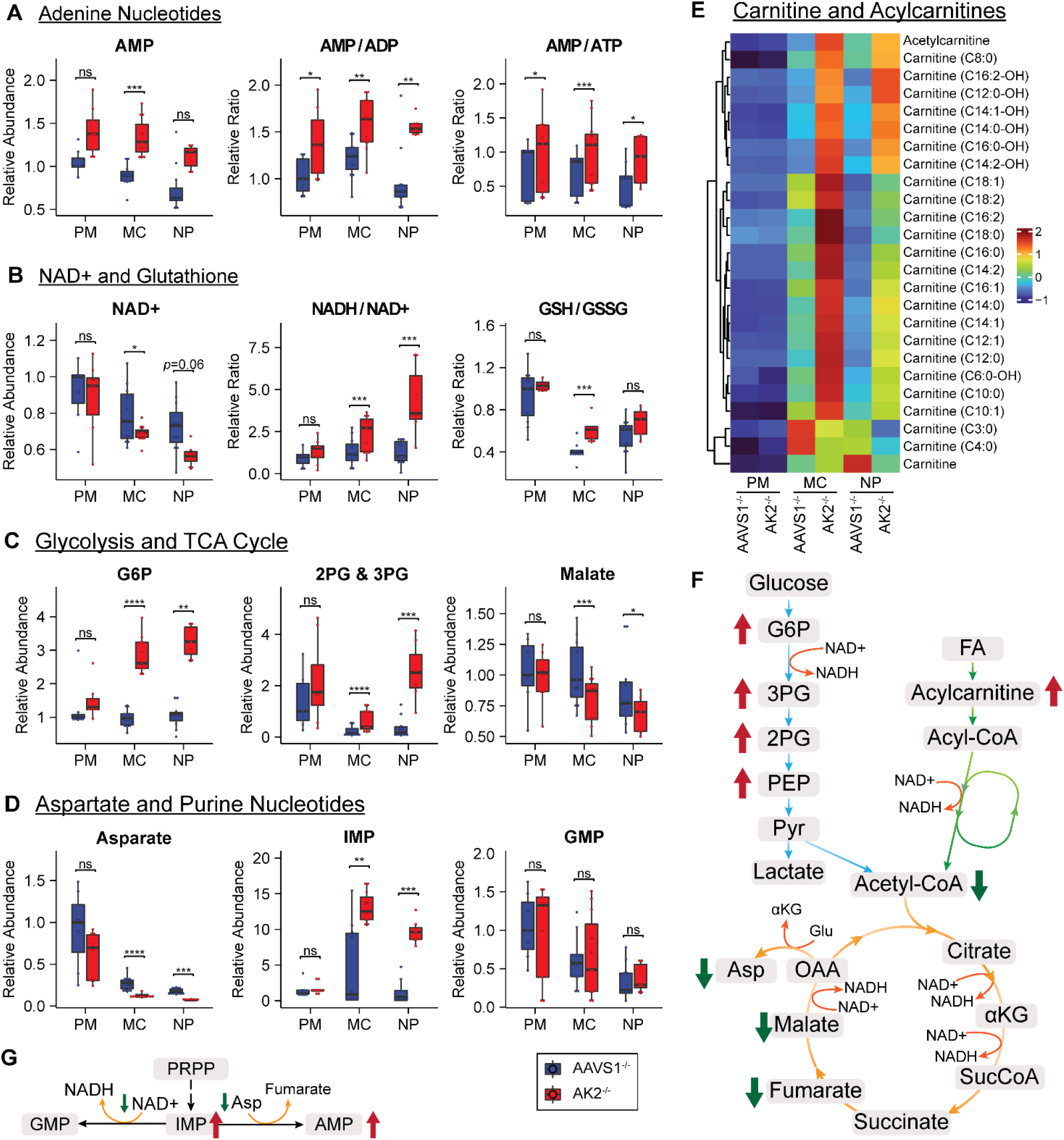
Metabolomics profiling reveals broadly decreased mitochondrial biosynthetic activity and increased levels of the ribonucleotide precursor IMP in AK2-deficient cells. (A-C) Box plots of relative abundance or relative ratios of metabolites in *AAVS1^-/-^* and *AK2^-/-^* PMs, MCs and NPs. GSH, reduced glutathione; GSSG, oxidized glutathione; G6P, glucose 6-phosphate; 2PG, 2-phosphoglycerate; 3PG, 3-phosphoglycerate. (D) Heatmap of carnitine and acylcarnitine relative abundance in *AAVS1^-/-^* and *AK2^-/-^* PMs, MCs and NPs. (E) Schematic map of energy metabolism pathways showing increased abundance of glycolysis metabolites and acylcarnitines, and decreased abundance of TCA cycle metabolites and aspartate in *AK2^-/-^* cells. PEP, phosphoenolpyruvate; Pyr, pyruvate; FA, fatty acid; αKG, α-ketoglutarate; SucCoA, succinyl-CoA; OAA, oxaloacetate; Asp, aspartate; Glu, glutamate. (F) Box plots of relative abundance of metabolites in *AAVS1^-/-^* and *AK2^-/-^* PMs, MCs and NPs. IMP, inosine monophosphate; GMP, guanosine monophosphate. (G) Schematic map of AMP and GMP synthesis from IMP. Synthesis of AMP requires aspartate, and synthesis of GMP requires NAD+. Up and down arrows indicate increased or decreased metabolite levels in *AK2^-/-^* cells relative to control. Metabolomics samples were collected from 2 independent experiments and 3-5 technical replicates from each experiment.

Furthermore, *AK2^-/-^* MCs and NPs showed a global shift in energy metabolism. We observed significant increases in the levels of glycolytic metabolites, including glucose 6-phosphate (G6P), 2-phosphoglycerate (2PG), 3-phosphoglycerate (3PG) and phosphoenolpyruvate (PEP), while a wide range of mitochondrial metabolites, including the TCA cycle intermediaries acetyl-CoA, fumarate and malate (Fig 5C) were significantly decreased. Especially, aspartate, synthesized from the TCA cycle intermediate oxaloacetate (OAA), showed a profound and progressive reduction in levels as the cells differentiated (Fig 5D). In addition, *AK2^-/-^* MCs and NPs exhibited an accumulation of acylcarnitines, a precursor for β-oxidation (Fig 5E). Accordingly, acylcarnitine accumulation may point to impaired β-oxidation^45–47^. These data suggest that *AK2^-/-^* MCs and NPs have an increase in glycolysis and a global decrease in mitochondrial metabolism (Fig 5F).

The metabolite with the greatest change between *AK2^-/-^* and *AAVS1^-/^*^-^, was the purine inosine monophosphate (IMP), with an over 30-fold increase in AK2-deficient neutrophils (Fig 5D). GMP levels were not affected by AK2 depletion (Fig 5D). The increase in IMP levels in AK2-deficient cells, combined with the depletion of NAD+ and aspartate, lead us to further investigate defects in purine ribonucleotide/ribonucleoprotein metabolism (Fig 5G).

### AK2 deficiency reduces cellular RNA content and ribosomal subunit expression

Purines are building blocks of DNA and RNA. Defects in purine metabolism are known causes of immunodeficiency syndromes with myeloid maturation defects^48, 49^. During myeloid maturation, cells undergo profound biosynthetic adaptations in RNA content and protein expression^17–21^, raising the possibility that the maturation and proliferation arrest in RD occurs at a time of increased biosynthetic demand. To explore this further, we quantified cellular RNA content, ribosome subunit expression and protein synthesis rate in AK2-deficient and control cells. During normal myelopoiesis, the total RNA content of control cells quantified by pyronin Y staining^50^ peaked at the MC stage (Fig 6A, top). *AK2^-/-^* MCs showed a profound reduction in RNA content as compared to *AAVS1^-/-^* control MCs (Fig 6A, bottom). As ribosomal RNA (rRNA) constitutes >85% of cellular RNA^51^, we next asked whether ribosome biogenesis is compromised by AK2 deficiency. During myeloid development, ribosomal subunit expression in control cells peaked at the promyelocyte stage and then slowly declined (Fig 6B, top). The majority of genes encoding ribosomal subunits showed a profound downregulation in *AK2^-/-^* PMs and MCs as compared to *AAVS1^-/-^* control cells (Fig 6B, bottom). To understand if the defect in purine metabolism and/or aspartate synthesis was pervasive enough to compromise protein expression, we quantified the protein synthesis rate based on O-propargyl-puromycin (OP-Puro) incorporation^52^. Protein synthesis peaked at the promyelocyte stage during normal myelopoiesis (Fig 6C, top), and AK2-deficient PMs showed a significant reduction in protein synthesis as compared to control PMs (Fig 6C, bottom). These findings suggest that AK2 deficiency limits RNA and protein biosynthesis during the most proliferative stages of myelopoiesis. These results are consistent with our single cell RNA-seq data derived from primary RD patient bone marrow cells in showing defects in RNA metabolism and ribonucleoprotein synthesis in RD.

**Figure 6.**
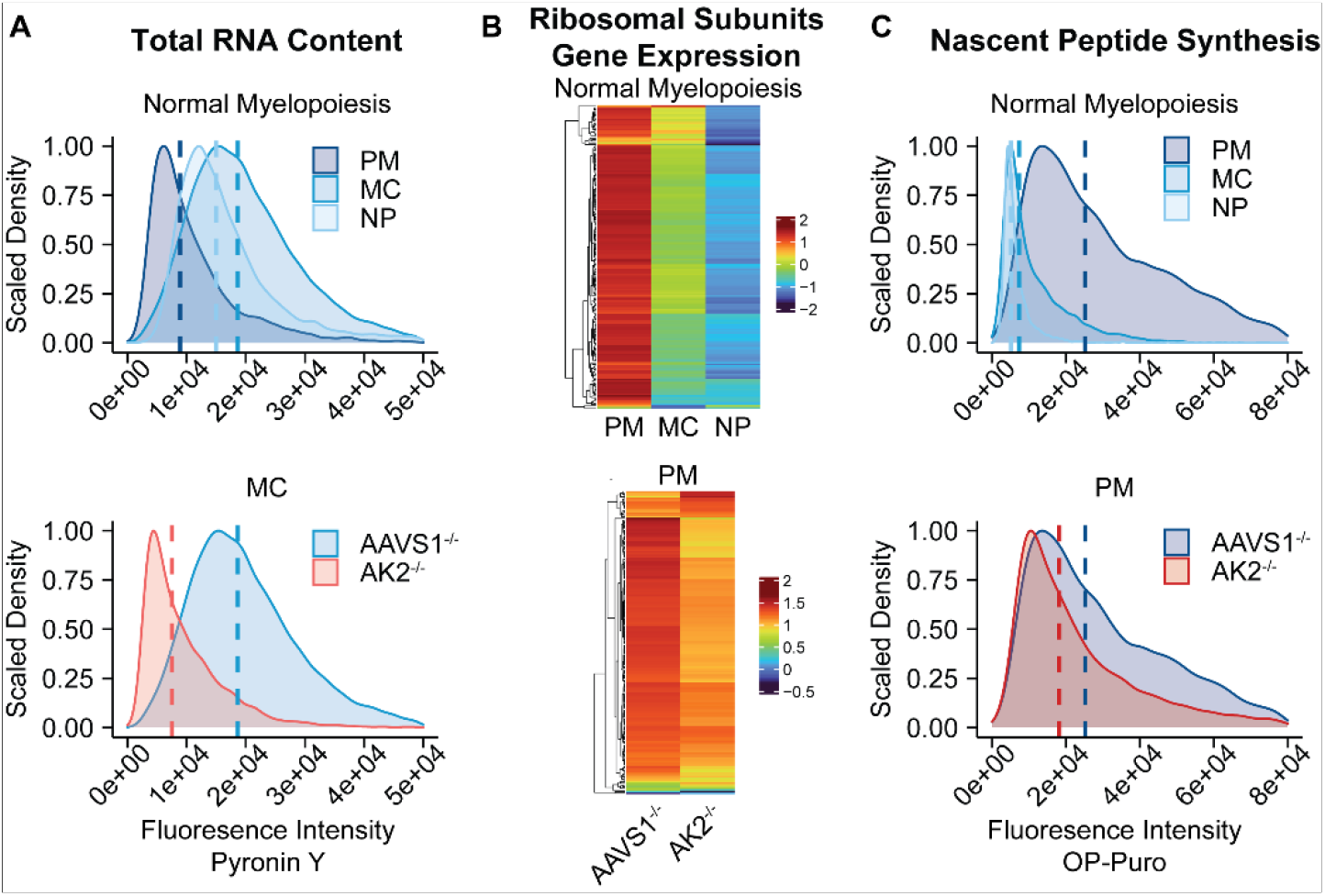
AK2 deficiency compromises cellular RNA content, ribosomal biogenesis and protein synthesis. (A) RNA content measured by pyronin Y staining. During normal myelopoiesis, RNA content peaked at the MC stage. *AK2^-/-^* MCs showed significantly lower RNA content than *AAVS1^-/-^* MCs. (B) Heatmaps of gene expression of ribosomal subunits, quantified by RNA-seq. During normal myelopoiesis, the expression of ribosomal subunit genes was highest at the PM stage. The majority of genes in this pathway were down regulated in *AK2^-/-^* PMs compared to *AAVS1^-/-^* control PMs. (C) Nascent peptide synthesis was measured by OP-Puro incorporation and staining. During normal myelopoiesis, peptide synthesis peaked at the PM stage. *AK2^-/-^* PMs showed significantly decreased peptide synthesis compared to *AAVS1^-/-^* PMs. For (A, C), representative flow cytometry results from at least 3 independent experiments were shown.

### NAD^+^ and aspartate deficiency do not confer purine auxotrophy in AK2-deficient cells

Next, we sought to investigate the nature of the defect in purine metabolism. Under normal physiological conditions, most of the cellular purine pool is derived from the recycling of degraded bases via the salvage pathway (Fig 7A, red)^53^. Under cellular conditions of increased purine demand, the *de novo* purine biosynthetic pathway is utilized (Fig 7A, green)^54–57^. The *de novo* pathway generates IMP from phosphoribosyl pyrophosphate, amino acids, formate and CO_2_. NAD^+^ and aspartate are critical for AMP and GMP biosynthesis from IMP^58^ (Fig 7A). Previous studies examining proliferation defects in respiratory chain-deficient cell lines found that NAD^+^ deficiency led to aspartate and purine auxotrophy^59, 60^. Purine auxotrophy and the resulting proliferation defects could be overcome in respiratory chain-deficient cells by restoring NAD^+^ and aspartate levels. Standard cell culture media contain purines in the form of nucleosides. To understand if NAD^+^ and aspartate depletion become limiting and confer purine auxotrophy during AK2-deficient myelopoiesis, we cultured *AK2*^-/-^ HSPCs in nucleoside-free culture medium and examined proliferation and differentiation. Any worsening of the RD phenotype from baseline upon culture in nucleoside-free medium would indicate that AK2-deficient cells are auxotrophic for purines. Surprisingly, AK2-deficient HSPCs grown in medium without nucleosides showed no aggravation of the RD phenotype but a trend towards improved proliferation and differentiation resulting in a slightly higher yield of MCs and NPs as compared to *AK2^-/-^* HSPCs grown in medium with nucleosides (Fig 7B). This raised the possibility that nucleosides were toxic to the cells rather than limiting due NAD^+^ and aspartate deficiency.

**Figure 7.**
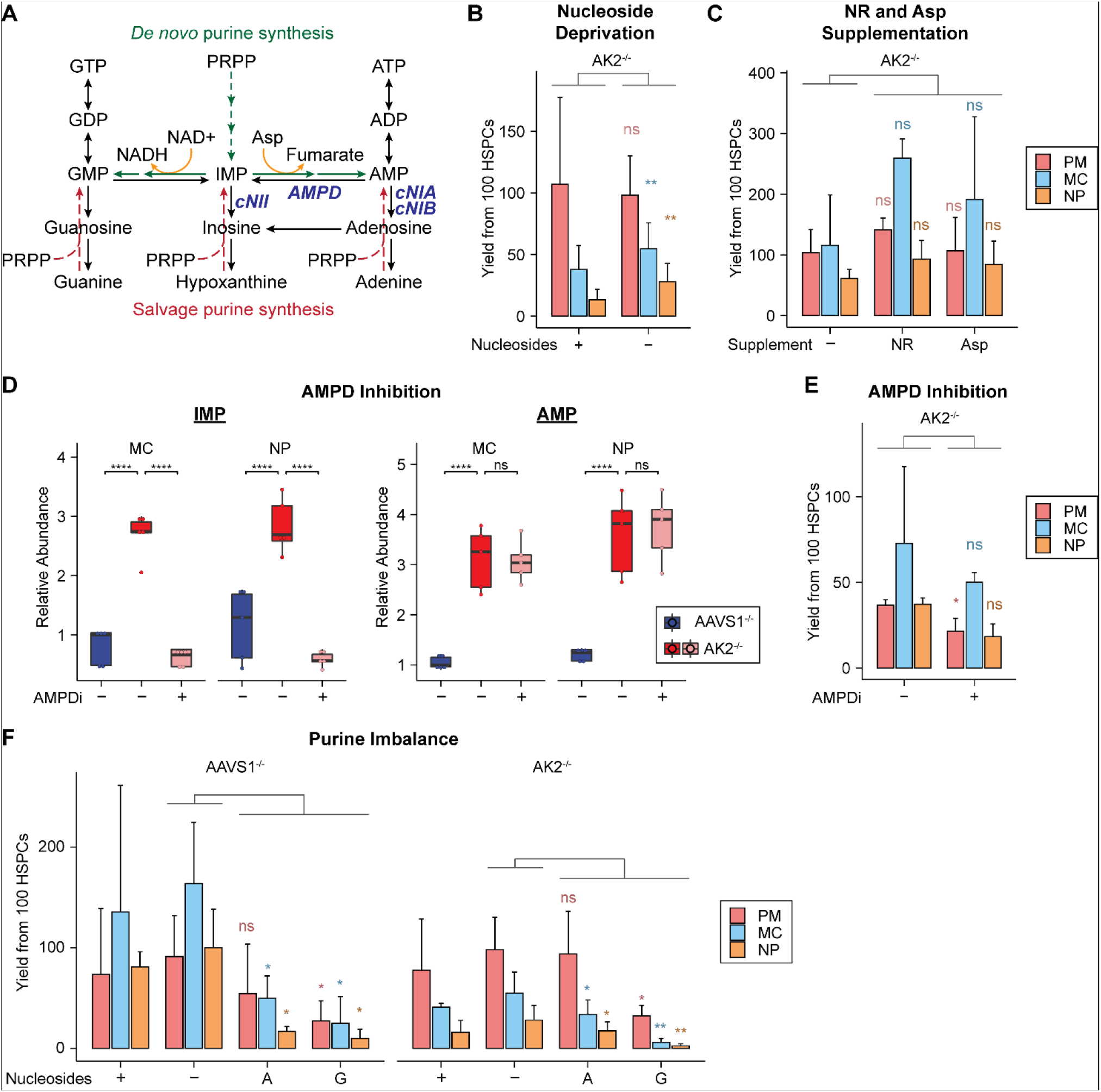
Increased AMP levels in AK2 deficiency disrupt purine metabolism. (A) Schematic map of purine biosynthesis, interconversion and degradation pathways. Purine nucleotides are synthesized *de novo* from PRPP (green), or salvaged from guanine, hypoxanthine and adenine (red). IMP serves as precursor for AMP and GMP synthesis. Aspartate and NAD+ are required for the synthesis of AMP and GMP from IMP, respectively. AMPD catalyzes the deamination of AMP to IMP (blue). Other important enzymes that degrade purine nucleotides include cytosolic 5’-nucleotidases (cN) IA, IB and II (blue). (B) Effects of nucleoside deprivation on myeloid development: Yield of *AK2^-/-^* PMs, MCs and NPs from 100 HSPCs differentiated in MEMα medium with (+) and without (-) nucleoside supplementation. (C) Effect of nicotinamide riboside (NR) and aspartate (Asp) supplementation on myeloid development: Yield of *AK2^-/-^* PMs, MCs and NPs from 100 HSPCs differentiated in Myelocult™ medium without (-) and with (+) supplementation of NR and Asp. (D, E) Effects of adenosine monophosphate deaminase (AMPD) inhibition on myeloid development: *AAVS1^-/-^* and *AK2^-/-^* cells were differentiated in Myelocult™ medium treated with (+) and without (-) a pharmacologic AMPD inhibitor (AMPDi). (D) LC-MS/MS quantification of IMP and AMP levels in *AAVS1^-/-^* and *AK2^-/-^* MCs and NPs with (+) and without (-) AMPDi treatment. Samples were collected from 2 independent experiments and 2-3 technical replicates from each experiment. (E) Yield of *AAVS1^-/-^* and *AK2^-/-^* PMs, MCs and NPs with (+) and without (-) AMPDi treatment from 100 HSPCs. (F) Effects of purine disequilibrium on myeloid development: Yield of *AAVS1^-/-^* and *AK2^-/-^* PMs, MCs and NPs from 100 HSPCs differentiated in MEMα medium with nucleosides (+), without nucleosides (-), with adenosine only supplementation (A) and with guanosine only supplementation (G). (B, C, E, F) showed average cell yield of 3 biological replicates. Error bars indicate standard deviation. Comparisons of yield are between treatment conditions as indicated, and between cells of the same differentiation stage.

To further explore this possibility, we tested several approaches to increase NAD^+^ biosynthesis or to alter the NADH/NAD^+^ ratio in the AK2-deficient HSPCs. We tried to boost NAD^+^ biosynthesis by supplementing with nicotinamide riboside (NR) (Fig 7C), nicotinamide, or nicotinic acid. We also tried to reduce NAD^+^ consumption by adding the sirtuin inhibitor FK866 or alter the redox balance by adding α-ketobutyrate, pyruvate, or oxidized glutathione (GSSG) (data not shown). Finally, we supplemented the cultures with aspartate (Fig 7C). None of these strategies rescued proliferation/myeloid differentiation in AK2-deficient HSPCs. Together, these data suggest that NAD^+^ and aspartate deficiency do not confer purine auxotrophy in RD.

### AK2-deficient cells deaminated excess AMP is to IMP

To test if high IMP levels in AK2-deficient cells are the result of increased AMP-breakdown, we pharmacologically inhibited the enzyme AMP-deaminase (AMPD), which converts AMP to IMP (Fig 7A, in blue) (AMPD inhibitor, sigma #5336420001)^61^. After AMPD inhibitor treatment, AK2-deficient cells exhibited a dramatic drop in IMP levels down to the level of control cells, while AMP levels in AK2-deficient cells underwent no significant change after AMPD inhibition (Fig 7D). The effect of AMPD inhibitor treatment suggests that high IMP levels in AK2-deficient cells result from deamination of AMP. The difference in fold-change between IMP and AMP levels after AMPD inhibition is at least partly reflective of the difference in pool size with the AMP pool being significantly larger than the IMP pool^62^. Although IMP levels in *AK2^-/-^* cells were rescued by AMPD inhibitor treatment, myelopoiesis did not improve, rather proliferation and differentiation showed a trend toward further decline(Fig 7E). These data raise the possibility that elevated AMP levels in AK2-deficient cells are toxic to myeloid progenitors.

### High AMP levels are toxic to granulocytes

To test if high AMP levels impair myelopoiesis, we induced a nucleotide disequilibrium by culturing *AK2^-/-^* and *AAVS1^-/-^* control HSPCs under four different conditions: (1) Culture in complete medium containing a balanced mix of nucleosides (including adenosine, deoxyadenosine, guanosine, deoxyguanosine, cytidine, deoxycytidine, thymidine and uridine); (2) culture in nucleoside-free medium; (3) and (4) culture in medium with addition of adenosine or guanosine, as the only added nucleoside source, respectively. The adenosine or guanosine concentration in adenosine-or guanosine-only medium was equal to the total purine nucleoside concentration in complete medium. In *AAVS1^-/-^* control HSPCs, culture in adenosine-or guanosine-only medium had catastrophic effects on proliferation and differentiation into MC and NP as compared to medium containing all or no nucleosides (Fig 7F). In contrast, *AK2^-/-^* HSPCs did not show any significant further deterioration from their already compromised baseline phenotype upon culture in adenosine-only medium. Culture in guanosine-only medium, however, further exacerbated the proliferation and differentiation defect of AK2-deficient cells (Fig 7F). Pyrimidine disequilibrium had no detrimental effects on either AK2-deficient or control cells (data not shown). The highly detrimental effect of adenosine-and guanosine-only culture on control cells supports the hypothesis that purine imbalance undermines myelopoiesis. The observation that adenosine-only culture medium has no additional detrimental effect on the maturation arrested phenotype of AK2-deficient cells compared to nucleoside-enriched or nucleoside-free culture, is consistent with the high endogenous AMP levels in AK2-deficient cells. Interestingly, while adenosine-and guanosine-only culture showed comparable results in control cells, the differential impact of guanosine-only culture on *AK2^-/-^* cells raises the possibility that endogenous AMP accumulation and exogenous guanosine disequilibrium have cumulative detrimental effects.

## DISCUSSION

Previous studies investigating the cellular consequences of Reticular Dysgenesis have examined the AK2 defect with focus on ADP depletion and its impact on OXPHOS-dependent ATP synthesis. Our study in highly purified *AK2^-/-^* human myeloid progenitor cells highlights the detrimental effects of AMP accumulation on purine metabolism. Increased intracellular AMP levels are known to control the *de novo* purine synthesis pathway by feedback inhibition of its rate limiting enzyme PRPP amidotransferase^61^.

Nonetheless, the effects of *AK2* deficiency are likely pleiotropic. We observed broad defects in mitochondrial metabolism and NAD^+^ and aspartate deficiency without a compromise in respiratory chain capacity. Although our studies did not support that NAD^+^ and aspartate deficiency confer purine auxotrophy upon purine deprivation, it is likely that NAD^+^ and aspartate depletion contribute on some level to the defects in ribonucleotide, ribonucleoprotein and protein biosynthesis.

We have considered the role of AMP-activated protein kinase (AMPK), which promotes catabolic processes and inhibits the mammalian target of rapamycin (mTOR) to suppress protein translation and cell growth^63 64, 65^; however, the levels of AMPK proteins in myeloid cells were very low and there was no significant difference in activated phospho-AMPK between *AK2^-/-^* and *AAVS1^-/-^* cells (Fig. S7), in line with the observation that AMPK deletion has little effect on adult hematopoiesis^66^.

Among the nine human adenylate kinase isoenzymes, only AK2 is located in the mitochondrial intermembrane space^6^. A number of studies have investigated the relationship between AK2 and cytosolic AK1. AK1 and AK2 catalytic activities are reversible and solely driven by surrounding substrate and product concentrations^6, 7^. Adenine nucleotides can permeate the outer mitochondrial membrane through voltage-dependent anion channels (VDACs). Conceptually, an increase in cytosolic AMP levels will eventually result in AMP phosphorylation to ADP by AK1. In support of this hypothesis, a recent report showed that *AK2* knockdown by short hairpin-RNA (shRNA) in primary T-ALL samples that did not express *AK1* led to apoptosis, while B-ALL samples with higher expression of *AK1* were less susceptible to *AK2* shRNA-knockdown^67^. However, other studies of *AK2* knockdown in HSPCs did not demonstrate rescue of hematopoiesis by ectopic *AK1* expression^13, 67^, raising the possibility that AK2 functions that are not redundant with AK1 are indispensable in cells that do not express *AK1*.

As AK isoenzymes with compensatory activity are not expressed in AK2-deficient myeloid cells, these cells may attempt to mitigate their AMP burden by degrading excess AMP by AMP deamination, leading to the observed increase in IMP. Although AMP deaminase-inhibitor treatment drastically reduced IMP levels in AK2-deficient cells, it did not significantly affect AMP levels. Considering the difference in pool size (AMP pool >>IMP pool)^62^, this suggests that AK2-deficient cells were only able to convert a small fraction of AMP into IMP. Whether a more efficient experimental reduction of AMP levels, e.g. through overexpression of AMP deaminase and/or 5’ cytosolic nucleotidases (Fig 7A, in blue) can rescue the proliferation and myelopoiesis defects in AK2-deficient cells, is subject of ongoing investigations.

Intriguingly, AMP and IMP catabolizing enzymes are involved in a wide range of hematopoietic disorders. Human AMP deaminase has at least 3 different isoforms (AMPD1/2/3), with AMPD2/3 being expressed in most hematopoietic lineages. Recently, loss of function mutations in *AMPD3* have been reported to cause T cell lymphopenia in mice^68^. Gain of function mutations in *NT5C2*, the gene encoding for cytosolic 5’ nucleotidase II, which converts IMP into inosine, are among the most frequently found mutations in relapsed ALL, present in 20% of relapsed T-ALLs and 3-10% of relapsed B-ALLs^69^, and are implicated in drug resistance to thiopurine based chemotherapy^69, 70^. These observations highlight the role tissue-specific expression patterns of purine metabolizing enzymes may play in defining characteristic, lineage-specific disease phenotypes.

We thus show that AK2 deficiency causes a defect in purine metabolism that leads to the accumulation of toxic levels of AMP, leading to defects in myelopoiesis. Understanding the distinct epistatic effects of purine metabolizing enzymes in different hematopoietic cell lineages may open-up new strategies to selectively target purine pathway networks specific to myeloid or lymphoid cell subsets to design the next generation of anti-proliferative agents with improved efficacy and decreased off target effects for immunosuppression and cancer therapy.

## Acknowledgements

KGW is a Uytengsu-Hamilton endowed faculty scholar at Stanford School of Medicine. SJM is a Howard Hughes Medical Institute (HHMI) Investigator, the Mary McDermott Cook Chair in Pediatric Genetics, the Kathryn and Gene Bishop Distinguished Chair in Pediatric Research, the director of the Hamon Laboratory for Stem Cells and Cancer, and a Cancer Prevention and Research Institute of Texas Scholar. AWD was supported by a Ruth L Kirschstein NRSA Fellowship. This work was also funded by the National Institutes of Health (AI123571 and DK11875). The BioHPC high performance computing cloud at UTSW was used for data analysis and storage as well as metabolomics data analysis (ODA) software deployment. Cord Blood Cells were made available by the Binns Program for Cord Blood Research. The Lorry Lokey Flow Cytometry Core was used for all flow cytometry.

---

**Figure S1.**
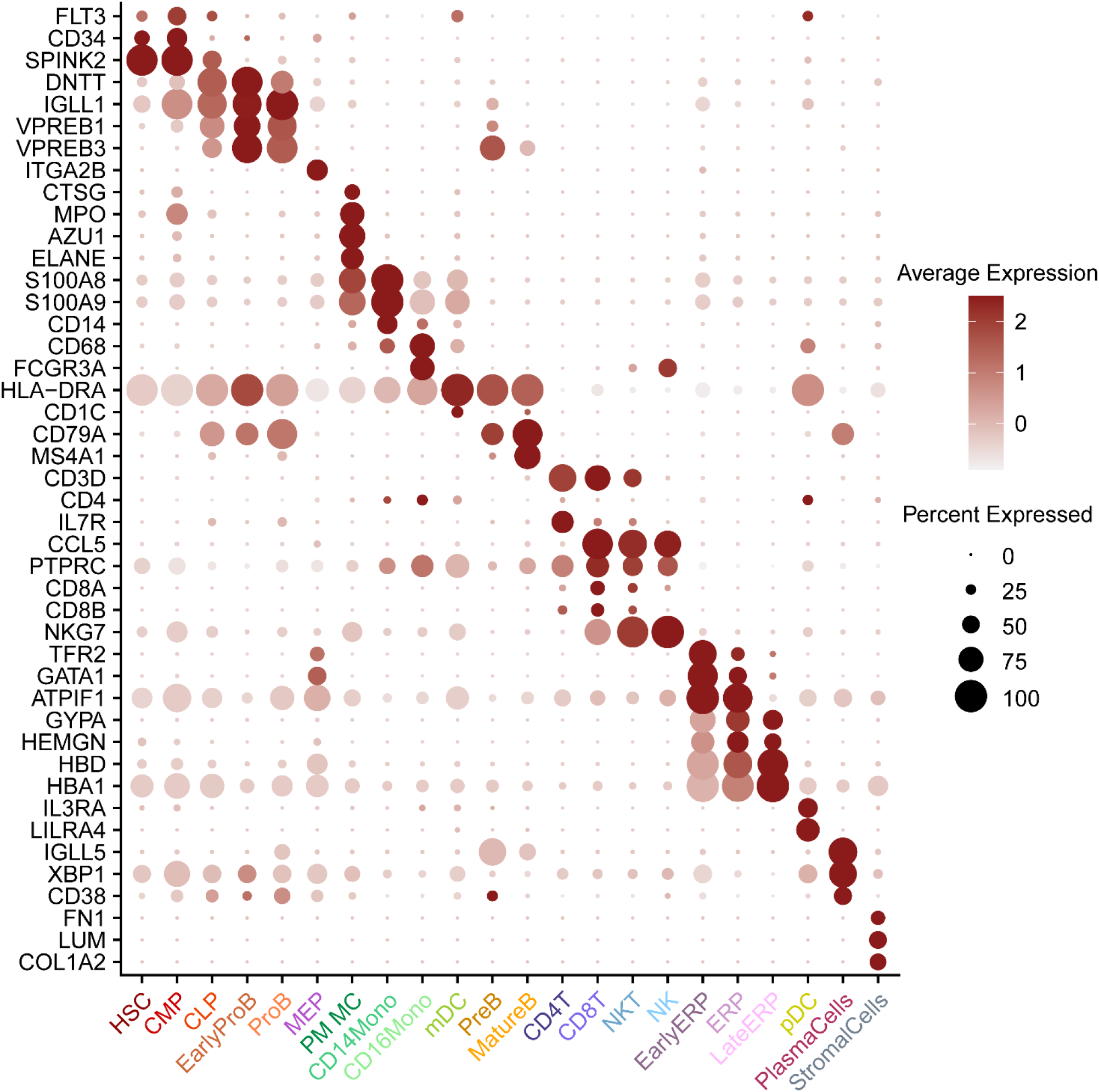
Gene expression of lineage specific markers in cell clusters identified in bone marrow scRNA-seq of RD patients and healthy donors. Dot size is proportional to the percentage of cells expressing a respective marker gene in an individual cluster; dot color intensity represents the average expression level of the respective marker gene in an individual cluster.

**Figure S2.**
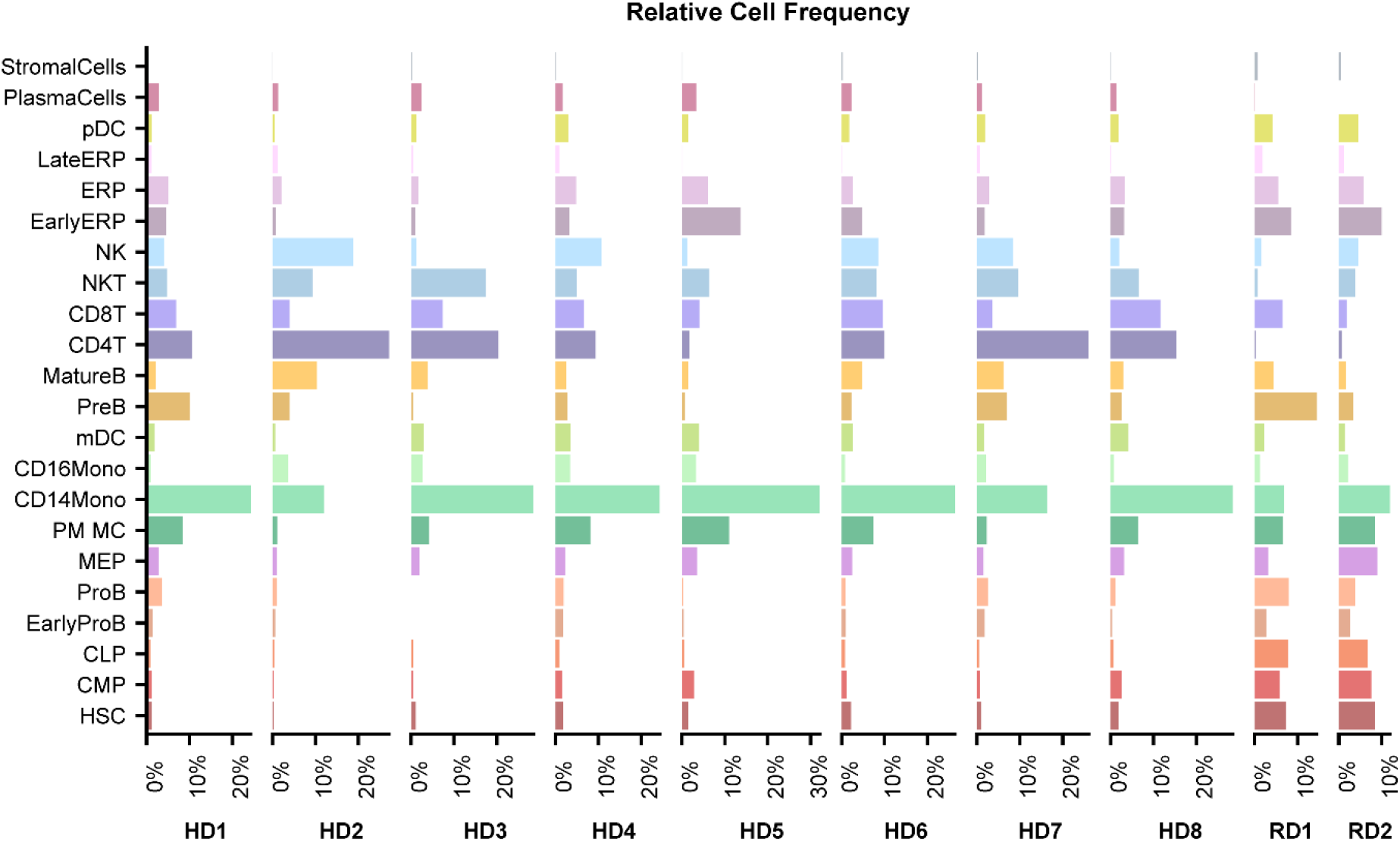
Relative frequency of cells per cluster identified by scRNA-seq in bone marrow of healthy donors and RD patients.

**Figure S3.**
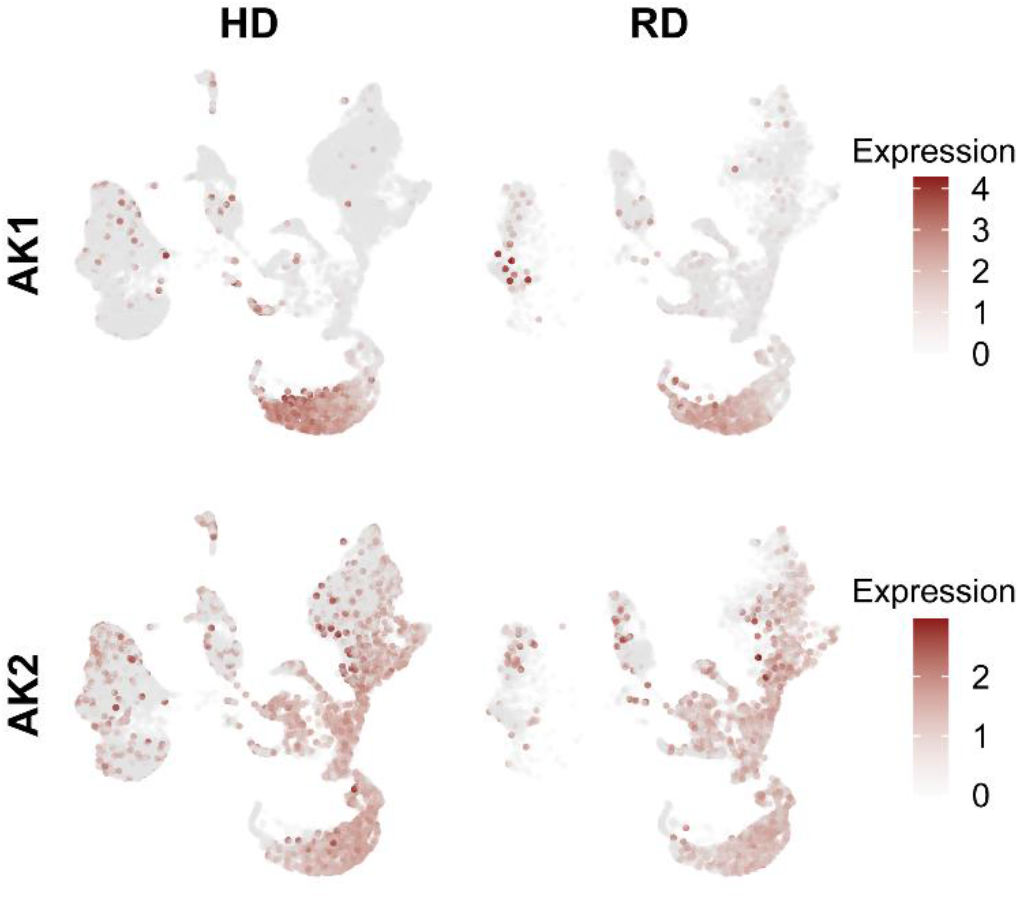
mRNA expression of *AK1* and *AK2* in bone marrow scRNA-seq of healthy donors and RD patients. *AK1* expression is largely limited in the erythroid lineage, and AK2 is broadly expressed in all hematopoietic lineages.

**Figure S4.**
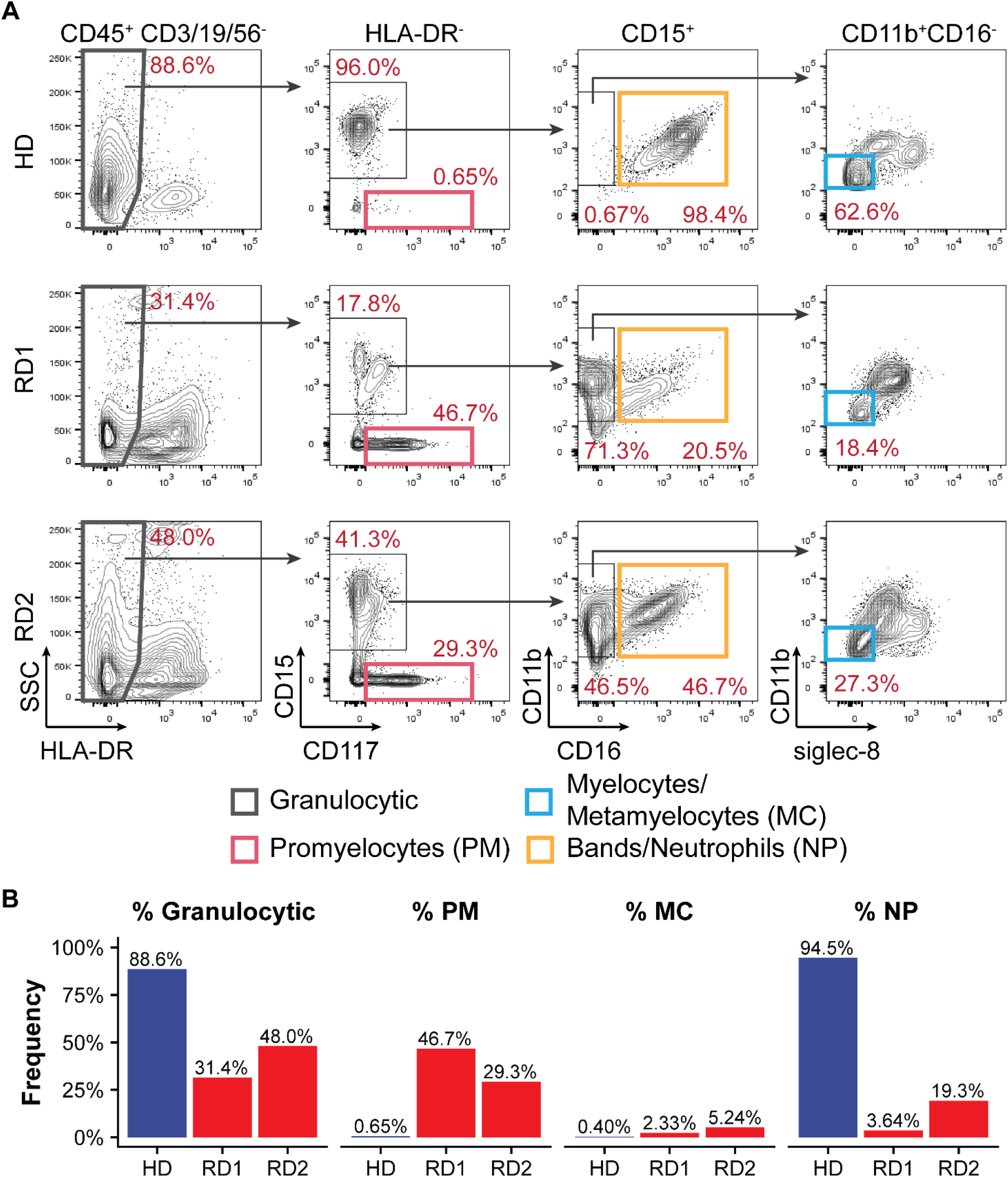
Flow cytometric analysis of RD patient bone marrow demonstrates myeloid maturation arrest at the promyelocyte stage. (A) Flow cytometric analysis of the myeloid lineage of healthy donor and RD patient whole bone marrow samples. Granulocytic lineage, CD45+ CD3-CD19-CD56-HLA-DR-; promyelocyte (PM), CD15+ CD117+; myelocyte/metamyelocyte (MC), CD15+ CD11b+ CD16-siglec8-; band/neutrophil (NP), CD15+ CD11b+ CD16+. (B) Bar plots showing percentages of granulocytic committed cells within the myeloid lineage, and percentages of PMs, MCs and NPs within the granulocytic lineage, in healthy donor and RD patients.

**Figure S5.**
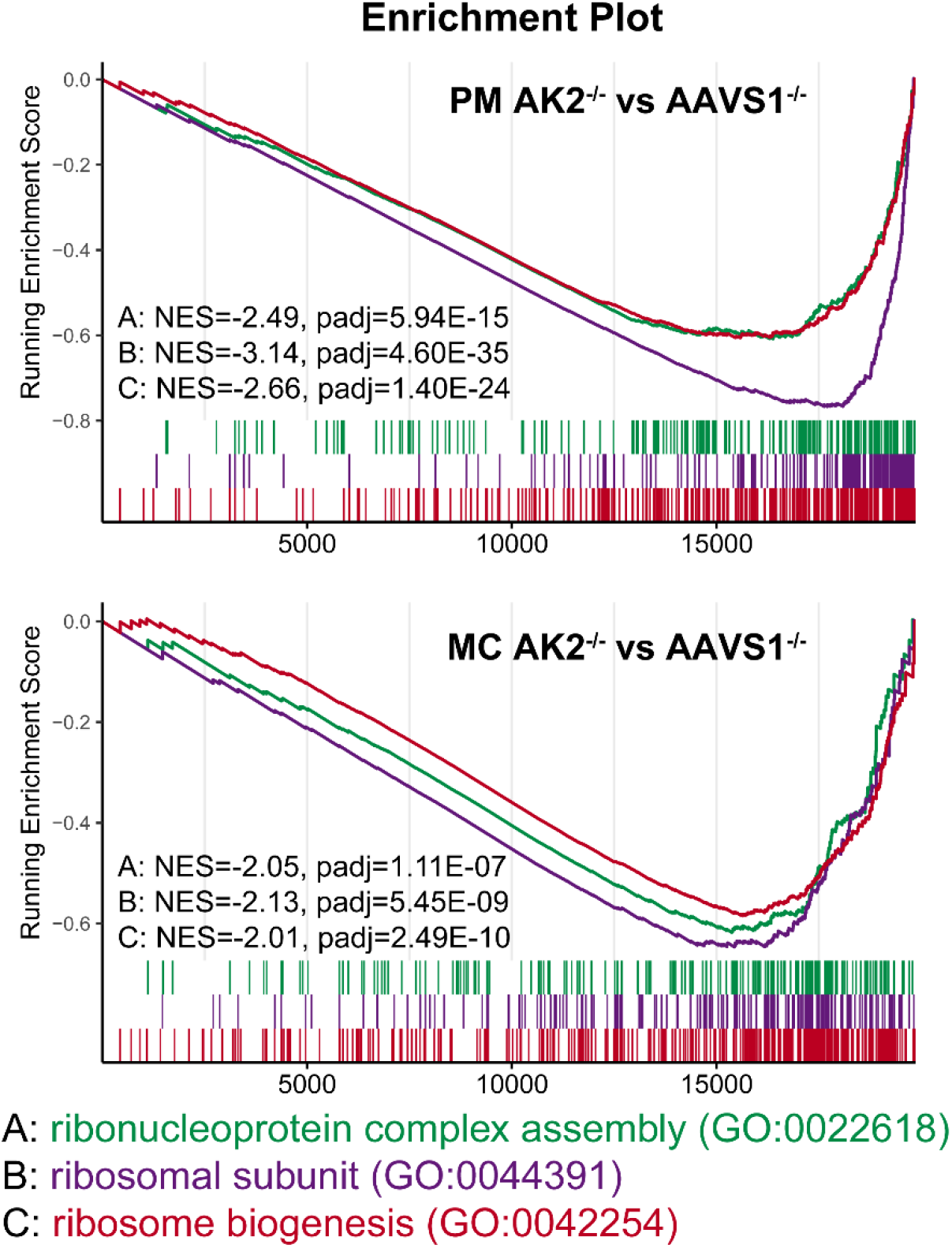
Gene set enrichment analysis of ribosome biogenesis related pathways comparing *AK2^-/-^* and *AAVS1^-/-^* PMs and MCs. Ribonucleoprotein complex assembly, ribosomal subunit expression and ribosome biogenesis pathways are significantly down regulated in *AK2^-/-^* cells at PM and MC stages, compared to *AAVS1^-/-^* cells at the same differentiation stage.

**Figure S6.**
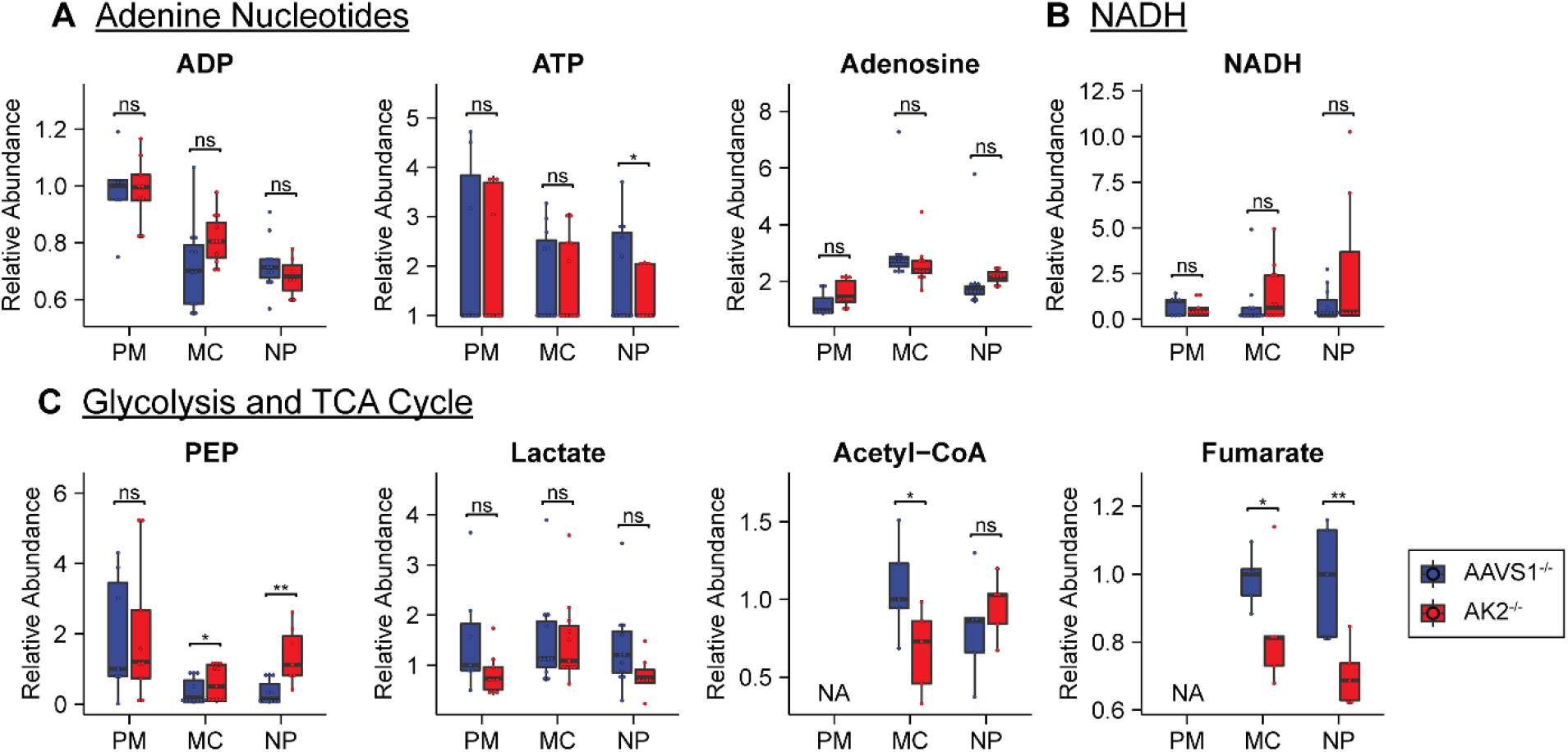
Selected metabolite levels by LC-MS/MS metabolomics in AK2-deficient and control cells. (A-C) Box plots of relative abundance or relative ratios of metabolites in *AAVS1^-/-^* and *AK2^-/-^* PMs, MCs and NPs. PEP, phosphoenolpyruvate.

**Fig S7.**
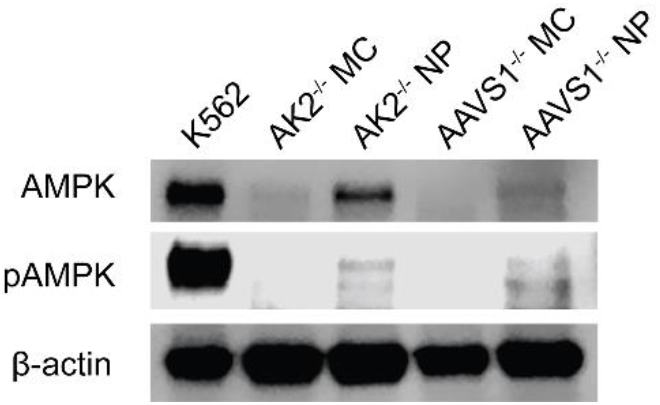
Western blot analysis of AMP-activated kinase (AMPK) and phospho (p)-AMPK in *AAVS1^-/-^* and *AK2^-/-^* cells. Total AMPK and pAMPK protein levels are low to undetectable in MCs. *AK2^-/-^* NPs exhibited higher AMPK levels than *AAVS1^-/-^* NPs, but comparable levels of pAMPK. K562 cells serve as a quality control for AMPK and pAMPK antibodies.

## REFERENCE

1. Lagresle-Peyrou, C. et al. Human adenylate kinase 2 deficiency causes a profound hematopoietic defect associated with sensorineural deafness. Nat. Genet. 41, 106–111 (2009).

2. Pannicke, U. et al. Reticular dysgenesis (aleukocytosis) is caused by mutations in the gene encoding mitochondrial adenylate kinase 2. Nat. Genet. 41, 101–105 (2009).

3. Hoenig, M., Pannicke, U., Gaspar, H. B. & Schwarz, K. Recent advances in understanding the pathogenesis and management of reticular dysgenesis. Br. J. Haematol. 180, 644–653 (2018).

4. Hoenig, M. et al. Reticular dysgenesis: international survey on clinical presentation, transplantation, and outcome. Blood 129, 2928–2938 (2017).

5. Vafai, S. B. & Mootha, V. K. Mitochondrial disorders as windows into an ancient organelle. Nature 491, 374–383 (2012).

6. Dzeja, P. & Terzic, A. Adenylate kinase and AMP signaling networks: metabolic monitoring, signal communication and body energy sensing. Int. J. Mol. Sci. 10, 1729–1772 (2009).

7. Dzeja, P. P. & Terzic, A. Phosphotransfer networks and cellular energetics. J. Exp. Biol. 206, 2039–2047 (2003).

8. Amiri, M., Conserva, F., Panayiotou, C., Karlsson, A. & Solaroli, N. The human adenylate kinase 9 is a nucleoside mono-and diphosphate kinase. Int. J. Biochem. Cell Biol. 45, 925–931 (2013).

9. Dzeja, P. P., Bast, P., Pucar, D., Wieringa, B. & Terzic, A. Defective metabolic signaling in adenylate kinase AK1 gene knock-out hearts compromises post-ischemic coronary reflow. J. Biol. Chem. 282, 31366–31372 (2007).

10. Janssen, E., Terzic, A., Wieringa, B. & Dzeja, P. P. Impaired intracellular energetic communication in muscles from creatine kinase and adenylate kinase (M-CK/AK1) double knock-out mice. J. Biol. Chem. 278, 30441–30449 (2003).

11. Henderson, L. A. et al. First reported case of Omenn syndrome in a patient with reticular dysgenesis. J. Allergy Clin. Immunol. 131, 1227–1230 (2013).

12. Rissone, A. et al. Reticular dysgenesis–associated AK2 protects hematopoietic stem and progenitor cell development from oxidative stress. J. Exp. Med. 212, 1185–1202 (2015).

13. Six, E. et al. AK2 deficiency compromises the mitochondrial energy metabolism required for differentiation of human neutrophil and lymphoid lineages. Cell Death Dis. 6, e1856 (2015).

14. Oshima, K. et al. Human AK2 links intracellular bioenergetic redistribution to the fate of hematopoietic progenitors. Biochem. Biophys. Res. Commun. 497, 719–725 (2018).

15. Chou, J. et al. Hypomorphic variants in AK2 reveal the contribution of mitochondrial function to B-cell activation. J. Allergy Clin. Immunol. 146, 192–202 (2020).

16. Geltink, R. I. K., Kyle, R. L. & Pearce, E. L. Unraveling the Complex Interplay between T Cell Metabolism and Function. Annual Review of Immunology 36, 461–488 (2018).

17. Skokowa, J., Dale, D. C., Touw, I. P., Zeidler, C. & Welte, K. Severe congenital neutropenias. Nat. Rev. Dis. Prim. 3, (2017).

18. Hoogendijk, A. J. et al. Dynamic Transcriptome-Proteome Correlation Networks Reveal Human Myeloid Differentiation and Neutrophil-Specific Programming. Cell Rep. 29, 2505–2519.e4 (2019).

19. Riffelmacher, T. et al. Autophagy-Dependent Generation of Free Fatty Acids Is Critical for Normal Neutrophil Differentiation. Immunity 47, 466–480.e5 (2017).

20. Cowland, J. B. & Borregaard, N. Granulopoiesis and granules of human neutrophils. Immunol. Rev. 273, 11–28 (2016).

21. Broxmeyer, H. E. et al. The importance of hypoxia and extra physiologic oxygen shock/stress for collection and processing of stem and progenitor cells to understand true physiology/pathology of these cells ex vivo. Current Opinion in Hematology 22, 273–278 (Curr Opin Hematol, 2015).

22. Makowski, L., Chaib, M. & Rathmell, J. C. Immunometabolism: From basic mechanisms to translation. Immunol. Rev. 295, 5–14 (2020).

23. Hay, S. B., Ferchen, K., Chetal, K., Grimes, H. L. & Salomonis, N. The Human Cell Atlas bone marrow single-cell interactive web portal. Exp. Hematol. 68, 51–61 (2018).

24. Stuart, T. et al. Comprehensive Integration of Single-Cell Data. Cell 177, 1888–1902.e21 (2019).

25. Bak, R. O., Dever, D. P. & Porteus, M. H. CRISPR/Cas9 genome editing in human hematopoietic stem cells. Nat. Protoc. 13, 358–376 (2018).

26. Aurnhammer, C. et al. Universal real-time PCR for the detection and quantification of adeno-associated virus serotype 2-derived inverted terminal repeat sequences. Hum. Gene Ther. Methods 23, 18–28 (2012).

27. Jha, P., Wang, X. & Auwerx, J. Analysis of Mitochondrial Respiratory Chain Supercomplexes Using Blue Native Polyacrylamide Gel Electrophoresis (BN-PAGE). Curr. Protoc. Mouse Biol. 6, 1–14 (2016).

28. Agathocleous, M. et al. Ascorbate regulates haematopoietic stem cell function and leukaemogenesis. Nature 549, 476–481 (2017).

29. DeVilbiss, A. W. et al. Metabolomic profiling of rare cell populations isolated by flow cytometry from tissues. Elife 10, 1–23 (2021).

30. Bray, N. L., Pimentel, H., Melsted, P. & Pachter, L. Near-optimal probabilistic RNA-seq quantification. Nat. Biotechnol. 34, 525–527 (2016).

31. Love, M. I., Huber, W. & Anders, S. Moderated estimation of fold change and dispersion for RNA-seq data with DESeq2. Genome Biol. 15, (2014).

32. He, L. et al. Detection and quantification of mitochondrial DNA deletions in individual cells by real-time PCR. Nucleic Acids Res. 30, (2002).

33. Grieshaber-Bouyer, R. et al. The neutrotime transcriptional signature defines a single continuum of neutrophils across biological compartments. Nat. Commun. 12, 1–21 (2021).

34. Zhang, S. et al. Adenylate kinase AK2 isoform integral in embryo and adult heart homeostasis. Biochem. Biophys. Res. Commun. 546, 59–64 (2021).

35. Waldmann, R. Institute for Transfusion Medicine Investigations on the Molecular Biology of Human Adenylate Kinase 2 Deficiency (Reticular Dysgenesis) and the Establishment and Characterisation of an Adenylate Kinase 2-deficient Mouse Model Dissertation Rebekka Waldm. (Universität Ulm, 2018). doi:10.18725/OPARU-6585

36. Patgiri, A. et al. An engineered enzyme that targets circulating lactate to alleviate intracellular NADH:NAD+ imbalance. Nat. Biotechnol. 38, 309–313 (2020).

37. Bak, R. O. et al. Multiplexed genetic engineering of human hematopoietic stem and progenitor cells using CRISPR/Cas9 and AAV6. Elife 6, 1–19 (2017).

38. Tiyaboonchai, A. et al. Utilization of the AAVS1 safe harbor locus for hematopoietic specific transgene expression and gene knockdown in human ES cells. Stem Cell Res. 12, 630–637 (2014).

39. Nakamura-Ishizu, A., Ito, K. & Suda, T. Hematopoietic Stem Cell Metabolism during Development and Aging. Dev. Cell 54, 239–255 (2020).

40. Chandel, N. S., Jasper, H., Ho, T. T. & Passegué, E. Metabolic regulation of stem cell function in tissue homeostasis and organismal ageing. Nat. Cell Biol. 18, 823–832 (2016).

41. Takubo, K. et al. Regulation of glycolysis by Pdk functions as a metabolic checkpoint for cell cycle quiescence in hematopoietic stem cells. Cell Stem Cell 12, 49–61 (2013).

42. Yu, W. M. et al. Metabolic regulation by the mitochondrial phosphatase PTPMT1 is required for hematopoietic stem cell differentiation. Cell Stem Cell 12, 62–74 (2013).

43. Loftus, R. M. & Finlay, D. K. Immunometabolism: Cellular metabolism turns immune regulator. J. Biol. Chem. 291, 1–10 (2016).

44. Al-Khami, A. A., Rodriguez, P. C. & Ochoa, A. C. Energy metabolic pathways control the fate and function of myeloid immune cells. J. Leukoc. Biol. 102, 369–380 (2017).

45. Rinaldo, P., Matern, D. & Bennett, M. J. Fatty acid oxidation disorders. Annu. Rev. Physiol. 64, 477–502 (2002).

46. McCoin, C. S., Knotts, T. A. & Adams, S. H. Acylcarnitines-old actors auditioning for new roles in metabolic physiology. Nat. Rev. Endocrinol. 11, 617–625 (2015).

47. Koves, T. R. et al. Mitochondrial Overload and Incomplete Fatty Acid Oxidation Contribute to Skeletal Muscle Insulin Resistance. Cell Metab. 7, 45–56 (2008).

48. Whitmore, K. V. & Gaspar, H. B. Adenosine deaminase deficiency - more than just an immunodeficiency. Front. Immunol. 7, (2016).

49. Grunebaum, E., Cohen, A. & Roifman, C. M. Recent advances in understanding and managing adenosine deaminase and purine nucleoside phosphorylase deficiencies. Curr. Opin. Allergy Clin. Immunol. 13, 630–638 (2013).

50. Rahmé, R. Assaying Cell Cycle Status Using Flow Cytometry. Methods Mol. Biol. 2267, 165–179 (2021).

51. Eddaoudi, A., Canning, S. L. & Kato, I. Cellular Quiescence. 1686, (Springer New York, 2018).

52. Hidalgo San Jose, L. & Signer, R. A. J. Cell-type-specific quantification of protein synthesis in vivo. Nat. Protoc. 14, 441–460 (2019).

53. Pedley, A. M. & Benkovic, S. J. A New View into the Regulation of Purine Metabolism: The Purinosome. Trends Biochem. Sci. 42, 141–154 (2017).

54. Henderson, J. F. & Khoo, M. K. Y. On the Mechanism of Feedback Inhibition of Purine Biosynthesis de Novo in Ehrlich Ascites Tumor Cells in Vitro. J. Biol. Chem. 240, 3104–3109 (1965).

55. Yamaoka, T. et al. Amidophosphoribosyltransferase limits the rate of cell growth-linked de novo purine biosynthesis in the presence of constant capacity of salvage purine biosynthesis. J. Biol. Chem. 272, 17719–17725 (1997).

56. Weber, G., Donohue, J. P., Glover, J. L., Natsumeda, Y. & Prajda, N. Enzymic Capacities of Purine de Novo and Salvage Pathways for Nucleotide Synthesis in Normal and Neoplastic Tissues. Cancer Res. 44, 2475–2479 (1984).

57. Mayer, D. et al. Expression of key enzymes of purine and pyrimidine metabolism in a hepatocyte-derived cell line at different phases of the growth cycle. J. Cancer Res. Clin. Oncol. 116, 251–258 (1990).

58. Salway, J. G. Metabolism at a Glance, 4th *Edition*. (Wiley-Blackwell, 2017).

59. Sullivan, L. B. et al. Supporting Aspartate Biosynthesis Is an Essential Function of Respiration in Proliferating Cells. Cell 162, 552–563 (2015).

60. Birsoy, K. et al. An Essential Role of the Mitochondrial Electron Transport Chain in Cell Proliferation Is to Enable Aspartate Synthesis. Cell 162, 540–551 (2015).

61. Admyre, T. et al. Inhibition of AMP deaminase activity does not improve glucose control in rodent models of insulin resistance or diabetes. Chem. Biol. 21, 1486–1496 (2014).

62. Zhao, H. et al. Quantitative analysis of purine nucleotides indicates that purinosomes increase de Novo purine biosynthesis. J. Biol. Chem. 290, 6705–6713 (2015).

63. Saito, Y. & Nakada, D. The Role of the Lkb1/AMPK Pathway in Hematopoietic Stem Cells and Leukemia. Crit. Rev. Oncog. 19, 383–397 (2014).

64. Hardie, D. G., Ross, F. A. & Hawley, S. A. AMPK: A nutrient and energy sensor that maintains energy homeostasis. Nat. Rev. Mol. Cell Biol. 13, 251–262 (2012).

65. Gowans, G. J., Hawley, S. A., Ross, F. A. & Hardie, D. G. AMP is a true physiological regulator of amp-activated protein kinase by both allosteric activation and enhancing net phosphorylation. Cell Metab. 18, 556–566 (2013).

66. Nakada, D., Saunders, T. L. & Morrison, S. J. Lkb1 regulates cell cycle and energy metabolism in haematopoietic stem cells. Nature 468, 653–658 (2010).

67. Maslah, N. et al. Adenylate kinase 2 expression and addiction in T-ALL. Blood Adv. 5, 700–710 (2021).

68. Zhan, X. et al. Adenosine monophosphate deaminase 3 null mutation causes reduction of naive T cells in mouse peripheral blood. Blood Adv. 4, 3594–3605 (2020).

69. Dieck, C. L. & Ferrando, A. Genetics and mechanisms of NT5C2-driven chemotherapy resistance in relapsed ALL. Blood 133, 2263–2268 (2019).

70. Moriyama, T. et al. Mechanisms of NT5C2-mediated thiopurine resistance in acute lymphoblastic leukemia. Mol. Cancer Ther. 18, 1887–1895 (2019).

71. Yu, G., Wang, L. G., Han, Y. & He, Q. Y. ClusterProfiler: An R package for comparing biological themes among gene clusters. Omi. A J. Integr. Biol. 16, 284–287 (2012).

